# A Female-Specific Microglial Redox Program Gates Susceptibility to Obesity

**DOI:** 10.64898/2026.09.04.749380

**Authors:** E Kyriakidou, G Cutugno, G Duranthon, P Gaikwad, E Archontoulaki, J Vergne, C Torrent, E Garbaye, S Lillo, O Kassem, D Jimenez-Blasco, P Zizzari, S James, V Simon, A Coutansais, N Dupuy, T Leste-Lasserre, G Laplagne, M David, J Ezan, F Martins, A Brochard, G Adisurja, B Salin, JA Reisz, G Marsicano, C Quarta, L Groc, UV Nägerl, JP Bolaños, R Rua, A D’Alessandro, A Mourier, C Allard, D Cota, A Nadjar

## Abstract

Chronic consumption of energy-dense, high-fat foods persistently exposes hypothalamic circuits that govern body weight to nutrient excess, progressively altering their activity and thereby promoting obesity. Microglia, the brain resident immune cells, sense circulating lipids, but how their intracellular metabolic programs adapt to chronic dietary excess, and how this contributes to obesity risk, is unclear. Here, we reveal pronounced sex differences in hypothalamic microglial responses to calorie overload. In females, but not males, microglia engage a protective metabolic program with increased antioxidant capacity and mitochondrial network remodeling, conferring resistance to early weight gain. Over time, activation of mTORC1 signaling disrupts mitochondrial function and dismantles this transient resilience, culminating in weight gain. These findings identify microglial mTORC1 as a sex-specific switch between resilience and vulnerability to obesity and position microglial metabolism as a tractable target for sex-informed weight control.

## Introduction

Chronic consumption of hypercaloric, fatty and ultra processed foods is a major driver of the contemporary obesity epidemic, promoting weight gain and metabolic disease^1–4^. As the central regulator of energy intake, expenditure and nutrient partitioning, the brain, and particularly the hypothalamus, is a key target of this energy excess, undergoing structural, functional and neuroimmune alterations in circuits that control body weight^5–10^. Elucidating how shifts in circulating energy substrates remodel the metabolic programs and activity patterns of brain cells may offer a critical opportunity to uncover novel mechanisms and therapeutic venues for the prevention and treatment of obesity.

Microglia are resident immune cells of the brain, where they monitor the environment, remove debris, and help control inflammation and repair^11–14^. Overnutrition and sustained excess caloric intake have been consistently linked to brain neuroinflammation, with microglia emerging as central intermediaries between diet induced inflammatory processes, altered neuronal function and increased susceptibility to weight gain^7,8,15^. In rodent models of high fat diet (HFD)-induced obesity, hypothalamic microglia rapidly proliferate and adopt a pro inflammatory phenotype^15–18^. Genetic or pharmacological dampening of microglial inflammatory signaling markedly reduces hyperphagia, weight gain and hypothalamic dysfunction, underscoring their bridge role between energy excess and metabolic imbalance^16,18–23^. Recent work has also identified hypothalamic microglia as nutrient and lipid sensors that directly respond to dietary saturated fatty acids and other circulating lipids, positioning these cells as dynamic integrators of metabolic cues within key neuronal circuits controlling whole body energy balance^17,24–26^. Microglia further undergo profound metabolic reprogramming upon injury or inflammatory stimulation, dynamically remodeling glycolysis, oxidative phosphorylation, and fatty acid oxidation through the engagement of signaling cascades such as the mechanistic target of rapamycin complex 1 (mTORC1) pathway^27–34^. This intracellular metabolic remodeling directly shapes microglial activation states, phagocytic activity, cytokine production, and overall functional responses in the brain^27–34^. However, how rapid shifts in microglial intracellular metabolism in response to changing dietary substrates reprogram their effector functions, and in turn shape energy balance and long term obesity risk, remains largely unknown.

Our data reveal striking sex differences in how microglia adapt to HFD. In females, but not males, microglia initially mount a protective metabolic program, with enhanced antioxidant capacity, mitochondrial fragmentation and lipid droplet buildup, that coincides with resistance to initial weight gain. Over time, activation of mTORC1 signaling in microglia emerges as a key driver of weight gain through mitochondria dependent mechanisms, underscoring microglial mTORC1 as a critical sex-dependent switch between resilience and vulnerability to obesity.

## Results

### Female microglia rapidly engage antioxidant programs in response to calorie excess

To investigate the response of hypothalamic microglia to calorie overload, 8-week-old male and female C57BL/6 mice were fed a HFD for six weeks, with body weight and food intake monitored weekly. Males displayed significant weight gain relative to standard diet (“Chow”) fed controls by week 1, whereas females did so only by week 3 (Fig. 1A-B). Weekly food intake (in kcal) increased significantly in male mice at 1 week of HFD, compared to chow-fed control mice. In females, no significant difference between HFD and chow-fed group were observed (Fig. 1C). These results highlight the different responses in energy balance regulation of male and female mice when exposed to calorie excess^35^.

**Figure 1.**
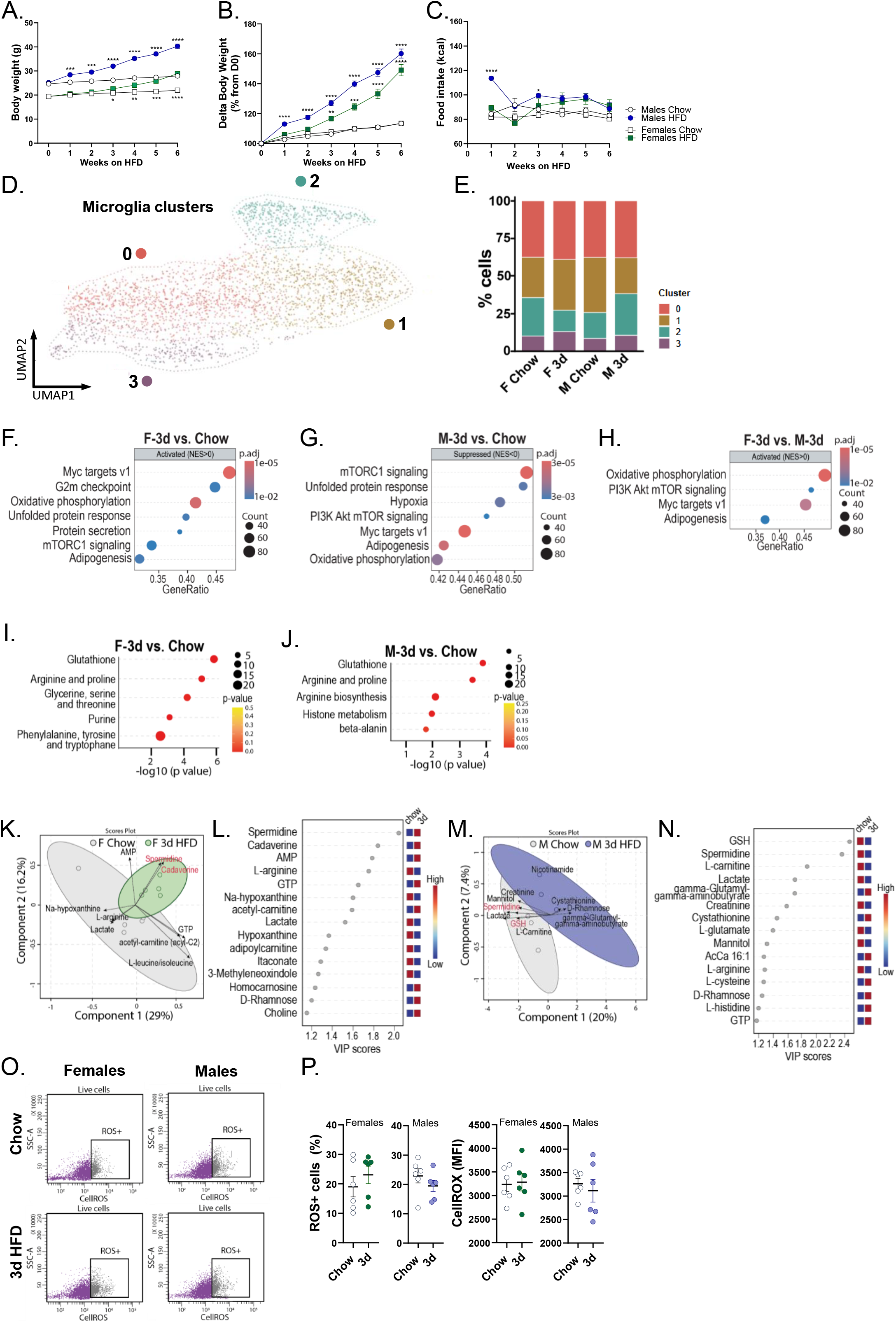
Acute lipid excess recruits anti-oxidant mechanisms in female microglia. **(A)** Weekly body weight measurements of female and male C57BL/6J mice fed a standard chow diet (Safe A03, 13.5% kcal from fat) or a high-fat diet (HFD, 60% kcal from fat) (n = 15, two-way repeated-measures [RM] ANOVA, *p<0.05, **p<0.01, ***p<0.001, or ****p <0.0001) **(B)** Body weight gain expressed as a percentage of body weight at the day of diet switch (n = 15, two-way RM ANOVA, **p<0.01, ***p<0.001, or ****p<0.0001 HFD versus Chow) **(C)** Weekly caloric intake following the diet switch (n = 15, two-way RM ANOVA, *p<0.05, or ****p<0.0001 HFD versus Chow) **(D)** UMAP embedding of hypothalamic microglia (clusters 0–3) from female and male mice fed control chow or HFD for 3 days (3d) or 6 weeks (6w), colored by Seurat cluster. **(E)** Stacked bar plot of microglial cluster proportions across the four experimental conditions. **(F-H)** Hallmark GSEA of pooled clusters 0–3, comparing female HFD 3d vs. female chow (F), male HFD 3d vs. male chow (G), and female vs. male at HFD 3d (H). Genes ranked by sign(log FC) × −log (p) from Seurat FindMarkers (Wilcoxon); enrichment by clusterProfiler::GSEA against MSigDB Mouse Hallmarks (minGSSize = 50). Top pathways by adjusted p-value (BH < 0.05) shown per comparison; dot size = leading-edge gene count, color = adjusted p-value. Only positively enriched pathways reached significance in F and H; only negatively enriched pathways in G. **(I–N)** Metabolomic profiling of *ex vivo*–isolated hypothalamic microglia from female (I, K– L) and male (J,M-N) mice exposed to chow or HFD for 3 days. **(I-J)** Pathway enrichment analyses identify significantly altered metabolic pathways in female (I) and male (J) microglia. PCA biplots (K,M) illustrate group separation and the metabolites contributing most strongly to discrimination between chow- and HFD-fed animals. Relative metabolite abundance is shown in panels (L,N). **(O–P)** Representative flow cytometry plots and quantification of intracellular ROS levels and the proportion of ROS-positive microglia measured by CellROX fluorescence after 3 days of HFD exposure in male and female mice (n = 6, unpaired t-test) (MFI=mean fluorescence intensity).

Previous studies have suggested rapid changes (within hours/days) in microglia within the arcuate nucleus of the hypothalamus (ARH)^8,16,17,22,23,36,37^. We first assessed microglial density, number and length of branches, and microglial volume in the ARH of male and female mice after 1 or 3 days of HFD and found no change in either sex in any of the parameters analyzed (Fig. S1A–F). Consistently, two-photon imaging did not reveal any significant modification in process motility in ARH microglia across timepoints (Fig. S1G– H).

We then used single-cell RNA sequencing (scRNAseq) to characterize microglial molecular responses to the diet, given their well-established capacity to display diverse, stimulus-dependent activation states^11^. CD11b^high^/CD45^low^ microglia were isolated using fluorescence-activated cell sorting (FACS)^38,39^, from the hypothalamus of female and male C57BL/6 mice after 3 days of HFD, alongside chow-fed controls. Unsupervised clustering analysis of the integrated dataset identified 4 microglial clusters (Fig. 1D). Clusters 0-2 comprised cells with a transcriptional signature reminiscent of homeostatic microglia, while cluster 3 expressed Disease Associated Microglia (DAM)-like genes^40–42^. The relative proportions of clusters 0 and 3 remained largely stable across conditions. In contrast, cluster 1 expanded in female mice after 3 days of HFD, but decreased in males under the same dietary condition. Cluster 2 showed the opposite pattern, increasing in males and decreasing in females in response to HFD (Fig. 1E). Hallmark Gene Set Enrichment Analysis (GSEA) of pooled clusters 0-3 (to maximize statistical power) indicated that, relative to chow, female microglia were enriched for proliferative (“G2M checkpoint”) and metabolically active programs (“MYC targets v1”, “oxidative phosphorylation”), along with increased “mTORC1 signaling” and lipid-associated pathways (“adipogenesis”) when exposed to HFD for 3d (Fig. 1F). In contrast, male microglia displayed suppression of these pathways (Fig. 1G). Direct comparison between female and male microglia under 3d HFD highlighted “oxidative phosphorylation” and “PI3K/Akt/mTOR signaling” as the main differentially regulated pathways (Fig. 1H). A lipid metabolism score (Seurat::AddModuleScore, REACTOME_METABOLISM_OF_LIPIDS gene set) revealed a sex-specific temporal response to HFD. Female microglia showed an early rise at 3 days, with the reverse dynamic in males (Fig. S2A). Untargeted lipidomics on MACS-sorted hypothalamic microglia, coupled to Partial Least Squares-Discriminant Analysis (PLS-DA) revealed sex- and diet-dependent lipidomic profiles that shifted markedly after acute caloric excess (Fig. S2B), with class-level differences by sex and diet (Fig. S2C). Membrane phospholipid pathways were enriched in both sexes, but LION analysis showed a female-specific overrepresentation of triacylglycerol, lipid storage, and lipid droplet pathways (Fig. S2D) absent in males (Fig. S2E). Consistent with enhanced lipid accumulation, Plin2 immunostaining revealed a marked rise in lipid droplets in female microglia after 3 days of HFD, but not in males (Fig. S2F-G). These data show that lipid excess affected both male and female hypothalamic microglia, evidenced by a marked perturbation of their lipid metabolism. To further validate and quantify the scRNAseq programs, we performed bulk RNAseq on hypothalamic microglia, using an optimized MACS-based hypothalamic microglia^43^ (Fig. S2H). After 3 days on HFD, no genes survived FDR correction (FDR < 0.1) in either sex, and proteomic analyses likewise showed few significantly modulated proteins (Fig. S2I), indicating that transcriptional and protein changes are subtle at this early stage. However, 251 genes tended to be differentially expressed (p < 0.05, fold change ≥ 2) in females and 608 in males (Fig. S2I-J). Gene Ontology analysis confirmed our GSEA findings. Female microglia were enriched for metabolic pathways (carbohydrate derivative catabolism, amino acid metabolism, lipid biosynthesis), whereas male microglia were enriched for immune and cell division pathways (activation of immune response, humoral immunity, mitotic spindle organization) (Fig. S2K-L). Taken together, these findings suggest that female microglia respond to acute HFD exposure by mobilizing metabolic pathways, including mTORC1 signaling and mitochondrial pathways. Male microglia, in contrast, engage immune mechanisms, consistent with the existing literature^7,8,18,21–23^.

To follow up on these findings, we next conducted mass spectrometry-based profiling of the intracellular metabolome in *ex vivo*–isolated hypothalamic microglia. PLS-DA revealed sex-and diet-dependent metabolic profile of microglia. Pathway enrichment analysis identified glutathione metabolism as the pathway most strongly affected by HFD in both female and male microglia (Fig. 1I-J). To determine the direction of the modification and the metabolites involved, we examined the measured metabolite levels and VIP scores. This further analysis revealed that the two sexes responded in opposite directions. Female microglia accumulated the polyamines spermidine and cadaverine, and the PCA biplot confirmed that the vectors for these metabolites drove most of the separation between groups (Fig. 1K-L), suggesting a shift toward antioxidant capacity. Male microglia showed the opposite response, with reduced levels of the antioxidant metabolites glutathione (GSH) and spermidine, as reflected in the biplot and VIP-score heatmap (Fig. 1M-N). Thus, although glutathione metabolism emerged as the top-ranked pathway in both sexes, the underlying metabolite shifts diverged, indicating an antioxidant-biased profile in females versus glutathione depletion in males. To evaluate the functional consequences of these metabolic alterations, we quantified intracellular ROS production using CellROX, a redox-sensitive probe that fluoresces upon oxidation^44^. After 3 days of HFD, there was no difference neither in CellROX fluorescence intensity nor in the proportion of ROS-positive hypothalamic microglia in either male or female mice (Fig. 1O-P), suggesting that ROS homeostasis under acute lipid challenge is maintained through mechanisms beyond glutathione mobilization alone. These data also suggest that sex-divergent antioxidant reprogramming observed in females reflects a distinct metabolic strategy rather than a quantitatively superior capacity to suppress oxidative stress per se.

Altogether, our findings reveal that female and male hypothalamic microglia adopt fundamentally distinct metabolic programs in response to acute lipid excess. Female microglia engage a coordinated antioxidant reprogramming centered on glutathione biosynthesis and utilization, while male microglia show evidence of glutathione depletion, suggesting an altered oxidative response.

### Mitochondrial remodeling underpins the antioxidant program of female microglia

Given the central role of mitochondria in microglial metabolic reprogramming and antioxidant responses^45–47^, we next examined mitochondrial features in male and female ARH microglia under HFD. We used the CX3CR1^CreERT2^;MitoTag^Ki/+^ mice^48^, in which microglial mitochondria express a GFP-tagged epitope^49,50^, and assessed the mitochondrial network architecture (Fig. 2A-B). Total mitochondrial volume was unchanged in both sexes in response to acute HFD, despite a trend to increased mtDNA copy number (Fig. 2C-D). Moreover, the number of GFP-positive mitochondrial particles increased in female microglia across the 3 days of HFD, while it decreased in male microglia (Fig. 2E). Electron microscopy of ARH microglial mitochondria corroborated these morphological findings: total mitochondrial area and density were unchanged, but mitochondria from females were shorter (p = 0.012), with a trend toward reduced perimeter (p = 0.0668), whereas males showed no significant changes (Fig. 2F-J). Proteomic analysis of MACS-sorted microglia revealed modest diet-dependent changes in mitochondrial proteins within each sex, while sex-dependent differences were more pronounced, particularly across oxidative phosphorylation and mitochondrial transport/dynamics modules (Fig. 2K-L). We observed reduced expression of OXPHOS complex I-IV proteins in 3d HFD female microglia relative to chow while no differences were observed in male microglia (Fig. 2M). This suggests that the mitochondrial fragmentation observed in female microglia in response to 3d HFD reflects a sex-specific remodeling of mitochondrial morphology and potentially OXPHOS function required to control energy and redox homeostasis.

**Figure 2.**
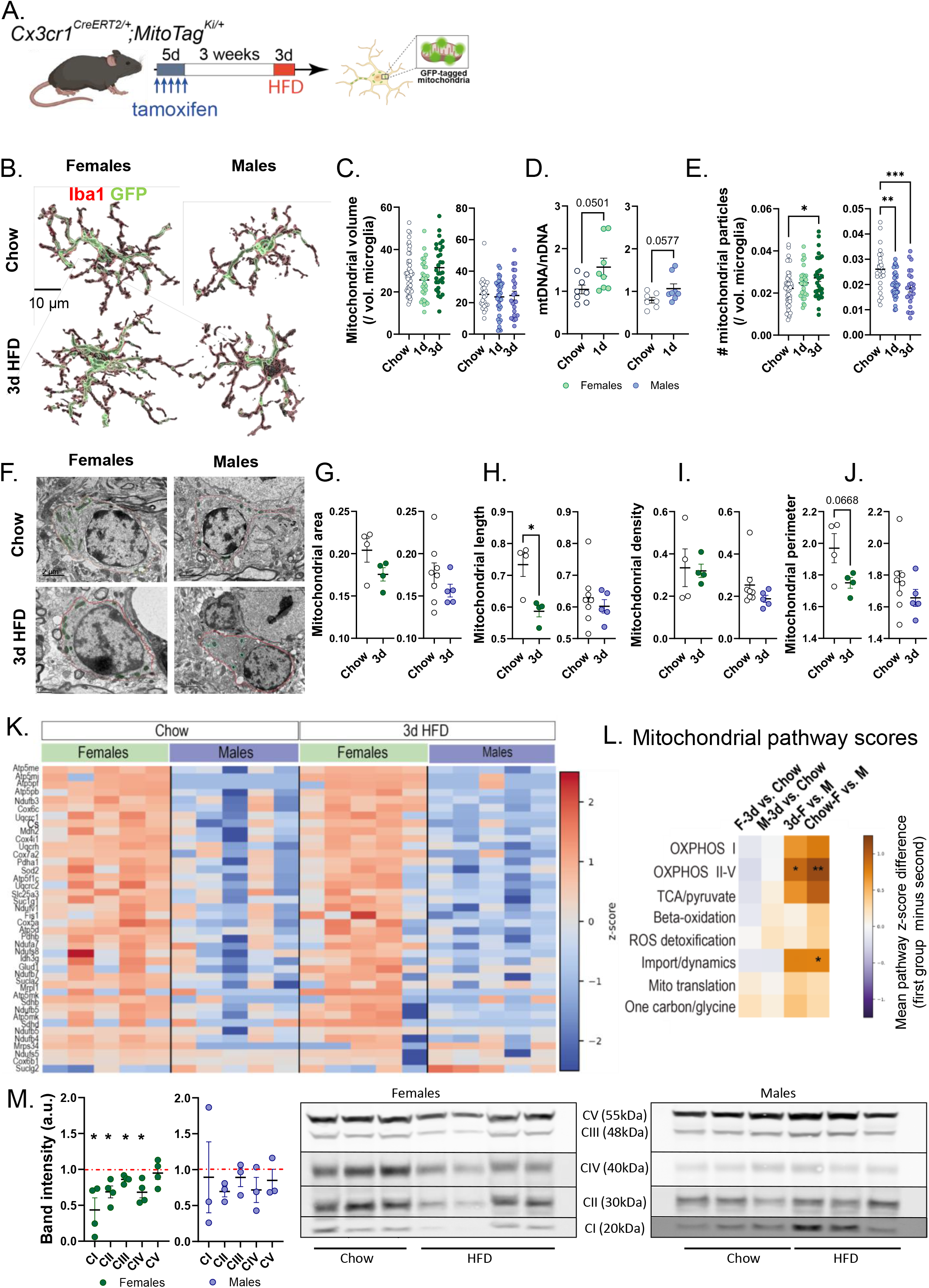
The female antioxidant program is built on mitochondrial remodeling under acute calorie excess. **(A)** Generation of the inducible microglia-specific mitoTAG (MG-MitoGFP) mouse model. Tamoxifen was administered to 8-week-old mice for 5 consecutive days and experiments were performed 3 weeks after the last injection to achieve selective labeling of microglial mitochondria. **(B)** Representative three-dimensional reconstructions of hypothalamic microglia immunostained for Iba1 and GFP in chow- and HFD-fed mice after 3 days of diet exposure. **(C)** Total mitochondrial GFP-positive particles volume per microglia following 1 or 3 days of HFD exposure. Each dot represents one cell (n = 26–45 cells from n = 5 mice/group, Kruskal-Wallis test). **(D)** Mitochondrial-to-nuclear DNA ratio (mtDNA/nDNA) in microglia isolated from mice fed a chow or HFD for 1 day (n=7-8, Welch’s test). **(E)** Total number of mitochondrial particles per microglial cell quantified from the reconstructions shown in (B). The same cells and number of animals were used as in (C) (n=26-45 microglia from n=5 mice/group, one-way ANOVA, *p<0.05, **p<0.01, ***p<0.001). **(F)** Representative electron microscopy images of hypothalamic microglia from chow- and HFD-fed mice after 3 days of diet exposure. Mitochondria are outlined in green and microglia in red. Scale bar = 2 µm. **(G–J)** Quantification of mitochondrial ultrastructural parameters, including mitochondrial area (G), length (H), density (I), and perimeter (J) (n=4-8, unpaired t-test, *p<0.05). **(K)** Heatmap showing z-scored abundance of mitochondrial proteins in female and male microglia under chow conditions and following 3 days of HFD exposure. **(L)** Mitochondrial pathway enrichment analysis showing pathway score shifts following 3 days of HFD exposure in female and male microglia. Positive values indicate pathway enrichment, whereas negative values indicate pathway depletion. **(M)** Quantification of the relative abundance of mitochondrial respiratory chain proteins in GFP-positive mitochondria isolated by immunoprecipitation following 3 days of HFD exposure (left panels; normalized to Ponceau S signal) and representative image of the western blot (right panels) (n=3-4, one sample t-test, *p<0.05).

Collectively, these findings indicate that female hypothalamic microglia meet acute lipid excess through a coordinated antioxidant program anchored in mitochondrial status, whereby remodeled mitochondria may sustain the redox defenses that counter lipid-induced oxidative stress, a sex-specific adaptation that possibly protects female microglia from oxidative damage and preserve brain function.

### Female microglial mTORC1 undermines antioxidant defenses under acute HFD

Single-cell RNA sequencing revealed the recruitment of mTORC1 pathway genes in hypothalamic microglia in response to 3d of HFD (Fig. 1F-H), a core signaling hub in hypothalamic energy sensing and food intake regulation^51^. Microglia rely extensively on kinase-driven signaling to sense and integrate immune and neuronal cues^52^. Therefore, to further elucidate the mechanisms underlying the divergent metabolic rewiring observed in male and female microglia in response to HFD, and to test for a potential implication of mTORC1 in this response, we conducted an unbiased functional kinome profiling of hypothalamic microglia.

As compared to chow, 3d HFD intake induced sex-specific and opposing signaling programs in microglia, with widespread increased kinase activity in females contrasted by a restricted and predominantly reduced signaling response in males, as observed on the kinome phylogenetic trees (Fig. 3A-B). PamGene kinase profiling further indicated that 3d HFD intake mobilizes several kinases in female microglia, including p70S6K, PKA, PKG, PKC, CaMK4, and Pim family kinases (Fig. S3A-B). These kinases are known to integrate nutrient- and energy-sensing signals with translational and survival pathways, confirming that microglia rapidly adjust their metabolic and signaling programs in response to lipid challenge. Among the top regulated kinases, and consistent with transcriptomic data (Fig. 1F-H), mTOR emerged as a key node, with activity increased in female but decreased in male microglia in response to HFD (Fig. 3A-B). Its downstream effector p70S6K was also significantly mobilized in female hypothalamic microglia only (Fig. S3A).

**Figure 3.**
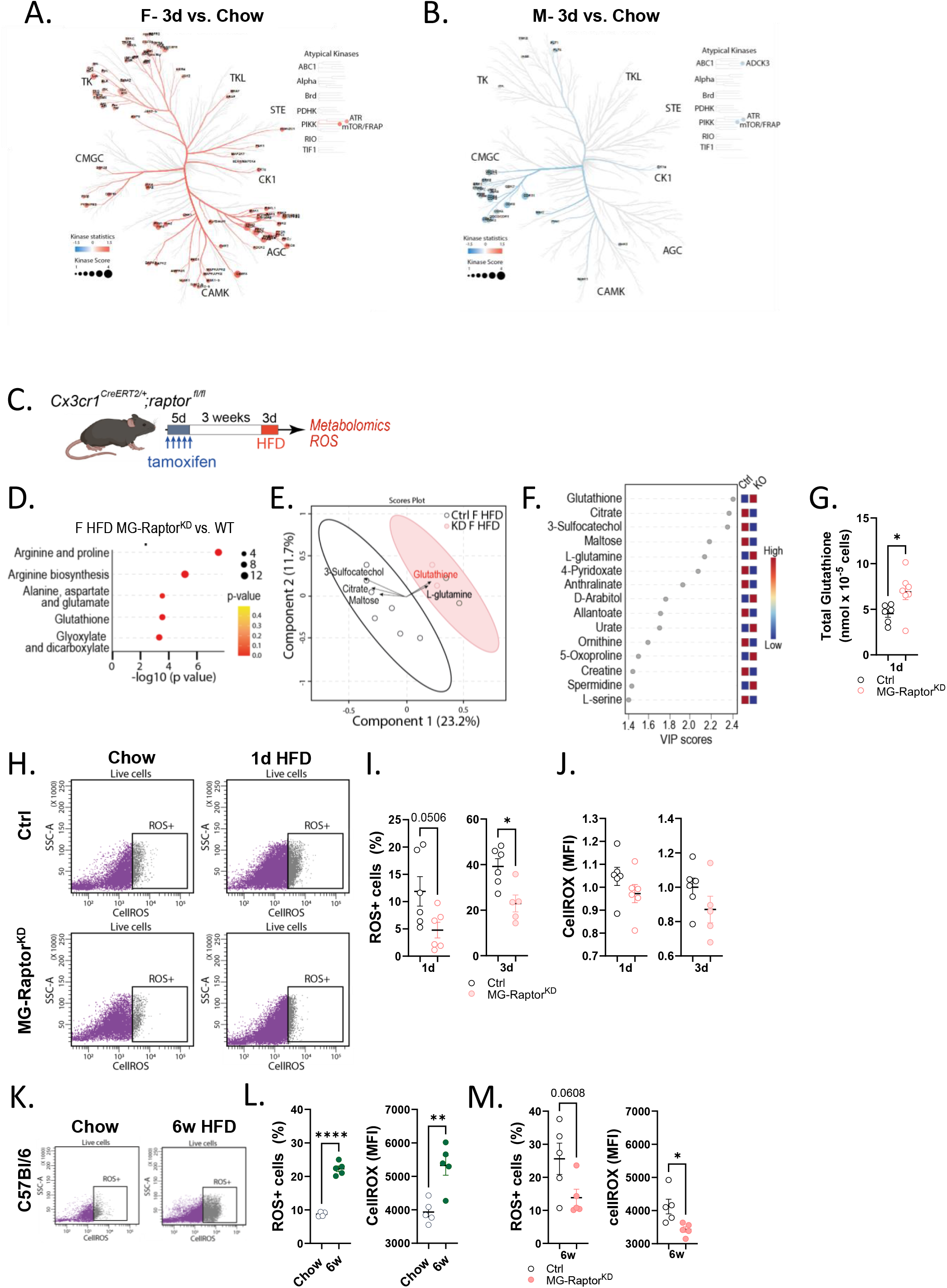
Acute mTORC1 signaling counteracts antioxidant responses in female microglia. **(A–B)** Kinome trees illustrating kinase activity profiles in female (A) and male (B) microglia following 3 days of HFD. Kinases are organized according to their phylogenetic families. Red lines indicate increased kinase activity, whereas blue lines indicate decreased kinase activity relative to chow-fed controls. Line thickness reflects the magnitude of the kinase activity change. **(C**) Generation of the inducible microglia-specific Raptor knock down (MG-Raptor^KD^) mouse model. Tamoxifen was administered to 8-week-old mice and experiments were performed 3 weeks after the last injection to achieve selective deletion of Raptor in microglia. **(D-F)** Metabolomic profiling of MACS-isolated hypothalamic microglia from HFD-fed control and MG-Raptor^KD^ female mice following 1 day of HFD exposure. Pathway enrichment analysis (D), PCA biplot (E), and VIP scores identifying metabolites contributing most strongly to group discrimination (F). In VIP plots, metabolites are ranked according to their contribution to group discrimination. Red indicates increased abundance and blue indicates decreased abundance in the indicated comparison. **(G)** Total glutathione levels in whole-brain microglia measured by enzymatic spectrophotometric assay in control and MG-Raptor^KD^ mice following 1 day of HFD exposure (n=6-7, Welch’s test, *p<0.05). **(H-J)** Representative flow cytometry plots and quantification of ROS-positive whole-brain microglia from control and MG-Raptor^KD^ mice measured by CellROX fluorescence after 1 day, and 3 days of HFD exposure (n=5-6, unpaired t-test, *p<0.05). **(K-L)** Representative flow cytometry plots and quantification of intracellular ROS levels and the proportion of ROS-positive microglia from C57BL/6 mice measured by CellROX fluorescence after 6 weeks of HFD exposure in female mice (n=4-5, unpaired t-test, **p<0.01, ****p<0.0001). **(M)** Quantification of ROS-positive whole-brain microglia and intracellular ROS levels from control and MG-Raptor^KD^ mice measured by CellROX fluorescence after 6 weeks of HFD exposure (n=5, unpaired t-test, *p<0.05).

Given that female microglia engage mitochondrial and antioxidant processes (Fig. 1-2) and the known link between mTORC1, mitochondrial function, and redox status^53–55^, we next investigated the role of this pathway in the response of female microglia to HFD. For this purpose, we generated CX3CR1^CreERT2^;Raptor^fl/fl^ mice, in which the *rptor* gene is conditionally knocked down specifically within microglia (MG-Raptor^KD^; Fig. S3C-F). Raptor is an essential component of mTORC1, acting as a scaffold that allows the kinase to phosphorylate metabolic effectors such as p70S6K^56,57^. Conditional deletion was confirmed to be brain-restricted and microglia-specific, with 83% excision efficiency in CD11b+ cells and no significant change in *rptor* expression in peripheral tissues or CD11b-brain cells (Fig. S3C-F). To then determine the contribution of mTORC1 signaling to the microglial kinome response, we profiled microglia from control (Ctrl) and MG-Raptor^KD^ female mice after 1 d HFD to capture the most early cellular changes. In Ctrl microglia, 1 d HFD triggered widespread kinome remodeling, with coordinated increases and decreases in kinase activity, including activation of the mTORC1 effector p70S6K, revealing again the rapid and dynamic kinome response to HFD, including mTORC1 cascade recruitment (Fig. S4A,C). By contrast, this response was largely absent in MG-Raptor^KD^ microglia under HFD (Fig. S4B,D), indicating that the kinome changes observed in Ctrl cells are largely dependent on mTORC1 signaling activation.

RNAseq analysis of MG-Raptor^KD^ hypothalamic microglia further revealed enrichment of gene ontology terms linked to cell division alongside changes in lipid metabolic pathways (Fig. S4E). However, despite this mitotic signature, quantification of microglial density showed no difference in cell number within the ARH across early time points (Fig. S4F), indicating that these transcriptional changes likely do not result in overt changes in proliferation acutely. To finally assess the effect of knocking down microglial Raptor on the neuronal activity within the hypothalamus, we used a multielectrode array (MEA), which allows real-time recording of spontaneous electrical activity across neuronal networks in acute hypothalamic slices^58^. As compared to Ctrl littermates, MG-Raptor^KD^ mice had decreased neuronal activity in the ventromedial and dorsomedial hypothalamus, suggesting a possible role for microglial mTORC1 in the remodeling of synaptic circuits in response to acute HFD^59,60^ (Fig. S4G-H).

To assess the role of the mTORC1 pathway in the HFD-induced anti-oxidant reprogramming of microglia, we next characterized the intracellular metabolome in *ex vivo*–isolated microglia from Ctrl and MG-Raptor^KD^ mice (Fig. 3C–G). Pathway enrichment analysis identified glutathione metabolism as one of the most upregulated in female MG-Raptor^KD^ microglia (Fig. 3D). PLS-DA revealed a genotype-specific metabolic signature in microglia (Fig. 3E), with glutathione emerging as a primary contributor to group separation (Fig. 3F). To corroborate these results, we quantified glutathione using a glutathione reductase–coupled enzymatic assay^61,62^ and observed elevated levels in female MG-Raptor^KD^ microglia after 1 □ d HFD (Fig. 3G), which was accompanied by a reduction in the proportion of ROS-positive microglia from 1 to 3 d HFD (Fig. 3H-J). We therefore asked whether the recruitment of the pro-oxidant mTORC1 pathway could drive a long-term increase in ROS production in C57BL/6 mice chronically fed an HFD. After 6 weeks of dietary exposure, female hypothalamic microglia showed a marked rise in both the proportion of ROS-positive cells and per-cell intracellular ROS levels (Fig. 3K-L), an increase abrogated in MG-Raptor^KD^ microglia exposed to chronic HFD (Fig. 3M), thus supporting a sustained inhibitory effect of mTORC1 activation on microglial anti-oxidant status.

Together, these data show that loss of mTORC1 signaling in female microglia recreates and prolongs the protective program normally seen in C57BL/6 females under HFD. Female microglia initially mount a robust antioxidant response, which is however maintained in time only when mTORC1 signaling is impaired. The resulting sustained redox shift identifies mTORC1 as a brake on the female antioxidant program whose removal locks microglia into a long lasting protective state.

### Microglial mTORC1 signaling promotes oxidation and body weight gain through mitochondrial reprogramming

To assess whether the enhanced and sustained antioxidant response of microglia following mTORC1 impairment leads to alterations in energy balance, mice were chronically exposed to HFD, with body weight and food intake monitored weekly over 6 weeks (Fig. 4A). HFD-fed MG-Raptor^KD^ female mice gained significantly less weight and had reduced fat mass as compared to Ctrl littermates (Fig. 4B-D), while lean mass and food intake were comparable between genotypes (Fig. 4D-E). Consistent with the kinome data (Fig. 3B), Raptor knockdown did not affect body weight gain in males (Fig. S4I).

**Figure 4.**
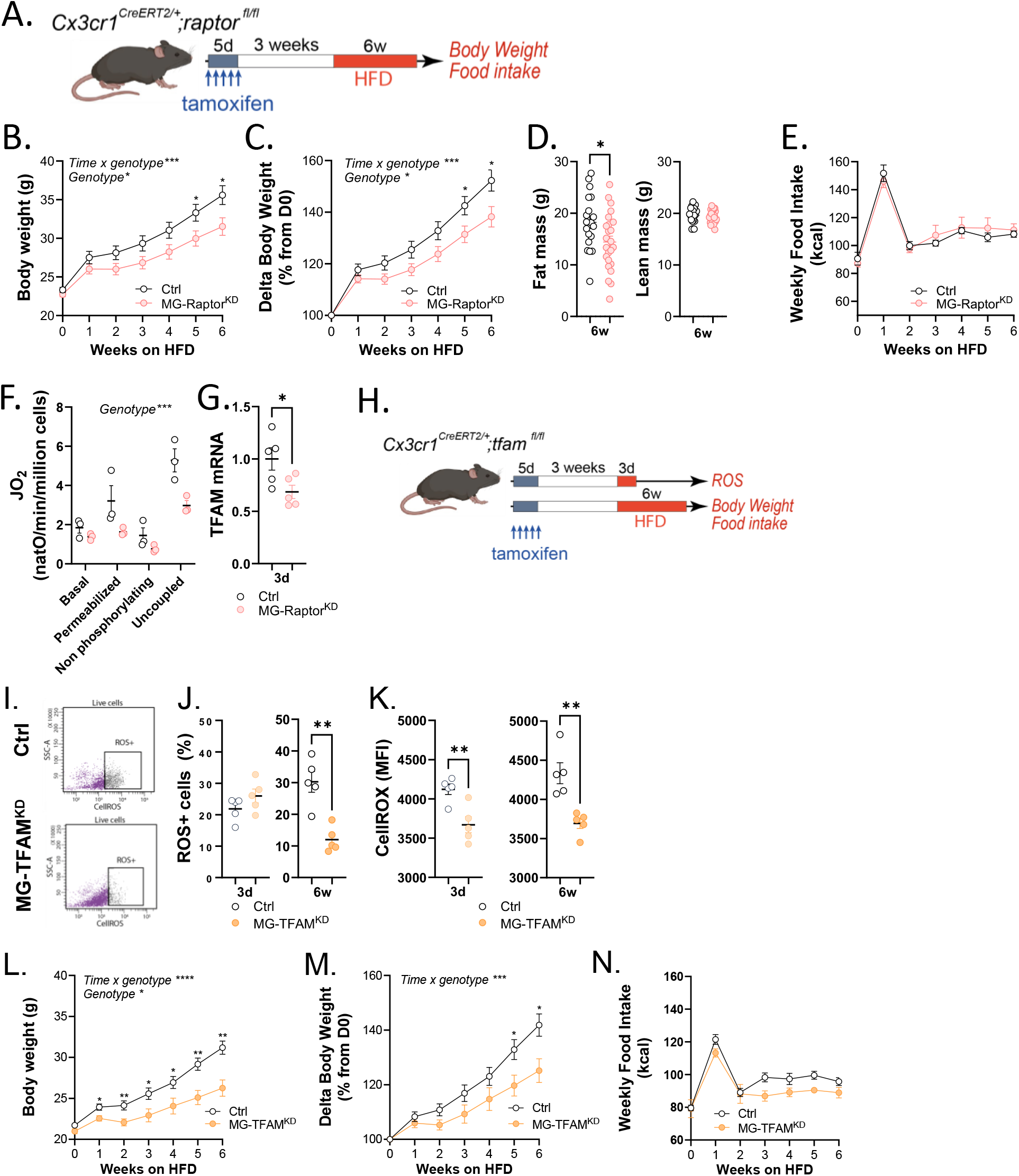
Microglial mTORC1 promotes oxidation and weight gain through mitochondrial reprogramming. **(A)** Experimental paradigm for long term assessment of body weight and food intake in MG-Raptor^KD^ mice. **(B)** Body weight (g) across the 6-week HFD regimen (n=19-24, 2 way-ANOVA RM, *p<0.05, ***p<0.001) **(C)** Percentage of body weight changes relative to baseline across the 6-week HFD regimen (n=19-24, 2 way-ANOVA RM, *p<0.05, ***p<0.001) **(D)** Fat and lean mass composition, expressed in grams (n=19-24, Unpaired t-test, *p<0.05). **E)** Averaged weekly food intake measurements expressed in kcal, of Ctrl and MG-Raptor^KD^ mice during chronic HFD exposure (n=19-24, 2 way-ANOVA RM). **(F)** Oxygen consumption rates of permeabilized microglia isolated from control and MG-Raptor^KD^ mice during the different respiratory states of the SUIT protocol. Values are expressed as oxygen flux (JO; amol O ·min ¹·cell ¹) (n=3, 2 way-ANOVA RM, ***p<0.001). **(G)** Relative *tfam* mRNA expression in microglia after 3 days of HFD exposure, normalized to control mice (n=5, unpaired t-test, *p<0.05). **(H)** Generation of the inducible microglia-specific *tfam* knock down (MG-TFAM^KD^) mouse model. Tamoxifen was administered to 8-week-old mice and experiments were performed 3 weeks after the last injection to achieve selective deletion of *Tfam* in microglia. **(I)** Representative flow cytometry plots showing CellROX fluorescence in microglia isolated from control and MG-TFAM^KD^ mice following HFD exposure. **(J-K)** Quantification of intracellular ROS levels measured by CellROX fluorescence intensity (J) and percentage of ROS-positive microglia (K) after 3 days and 6 weeks of HFD exposure in control and MG-TFAMKD mice (n=5, unpaired t-test, **p<0.01). **(L)** Body weight (g) across the 6-week HFD regimen (n=7-19, 2 way-ANOVA RM, *p<0.05, **p<0.01, ****p<0.0001) **(M)** Percentage of body weight changes relative to baseline across the 6-week HFD regimen (n=7-19, 2 way-ANOVA RM, *p<0.05, ****p<0.0001) **(N)** Averaged weekly food intake of Ctrl and MG-TFAM^KD^ mice during chronic HFD exposure (n=7-19, 2 way-ANOVA RM)

As a central regulator of cellular bioenergetics, mTORC1 governs both glycolytic flux and mitochondrial oxidative phosphorylation, each of which shapes cellular ROS balance, prompting us to examine both bioenergetic arms^63^. We first examined the role of HIF-1α, a key downstream effector through which mTORC1 promotes glycolysis and coordinates mitochondrial mass and function in immune cells, including microglia^64,65^. To this end, we quantified the microglial expression of HIF-1α transcripts in response to 3d or 6w of HFD and could not observe any difference between groups (Fig. S5A). We then employed CX3CR1^CreERT2^;HIF1α^fl/fl^ mice (‘MG-HIF1α^KD^’) in which the *hif1*_α_ gene is knocked-down specifically in microglia following tamoxifen treatment (Fig. S5B). After verifying the significant decrease in microglial HIF1α mRNA expression (Fig. S5C), we exposed female MG-HIF1α^KD^ and their Ctrl littermates to HFD for several weeks, but observed no genotype-dependent differences in body weight gain, neither in females (Fig. S5D), nor in males (Fig. S5E). We next focused on the mitochondrial arm of mTORC1-driven metabolic reprogramming. By using high-resolution respirometry (Oroboros), we found that mitochondrial respiration was significantly reduced in microglia from MG-Raptor^KD^ females as compared to Ctrl littermates (Fig. 4F), and associated with decreased mRNA expression of the mitochondrial transcription factor *tfam* (Fig. 4G). Building on this result, we next tested whether mTORC1 requires functional mitochondria to drive ROS production. To this aim, we employed CX3CR1^CreERT2^;TFAM^fl/fl^ mice (MG-TFAM^KD^) (Fig. 4H), leading to a significant decrease in the expression of *tfam* transcript in microglia (Fig.S5F). We confirmed that mitochondrial respiration was reduced in microglia while the total microglia number remained unaffected (Fig. S5G-H). We next evaluated microglial oxidative status and observed that the overall proportion of ROS-positive microglia was significantly reduced after 6w of HFD in female MG-TFAM^KD^ mice (Fig. 4I-J). Additionally, a significant reduction in microglial CellROX fluorescence intensity was observed in females after 3d and 6w of HFD (Fig. 4K), mirroring the decrease observed in female MG-Raptor^KD^ mice (Fig. 3H-J, M). Accordingly, MG-TFAM^KD^ females gained significantly less weight than their Ctrl littermates during chronic HFD despite similar food intake (Fig. 4L-N), an effect absent in males (Fig. S5I), thereby phenocopying the phenotype observed in female MG-Raptor^KD^ mice.

Altogether, these findings indicate that in female microglia, HFD rapidly activates an mTORC1-to-mitochondria axis that sustains oxidative metabolism and ROS production, and that attenuating this axis, either upstream at mTORC1 or downstream at the level of mitochondria, enhances the antioxidant tone of female microglia and uncouples HFD feeding from body weight gain.

## DISCUSSION

Hypothalamic microglia have been recognized as key lipid sensors, playing important roles in the development of obesity^7,8,39^. Our study now identifies microglia as essential regulators of energy homeostasis in the context of acute caloric excess, and shows that they govern this function in a sex-dependent manner. Rather than mounting the classical pro-inflammatory response seen in males^8,17,18,21–23,66,67^, female microglia adopt a metabolically adaptive state that coordinates mitochondrial network organization and antioxidant defenses. Early, concomitant engagement of mTORC1 signaling overwhelms these protective mechanisms, driving mitochondrial oxidative phosphorylation, increasing ROS production and ultimately shifting the balance toward weight gain during prolonged HFD intake. These data provide a mechanistic explanation for the female metabolic phenotype under HFD characterized by initial resistance followed by delayed weight gain, and position hypothalamic microglia at the core of this sexually dimorphic response.

One of the central findings of our study is that female hypothalamic microglia exhibit a distinct cellular metabolic state under HFD compared with males. Several factors are likely to contribute to this divergence. Previous work has shown that chronic HFD induces sex-specific remodeling of the brain lipid milieu, with males accumulating higher levels of saturated fatty acids (SFA) and sphingolipids in the central nervous system, whereas females maintain lower levels of these lipids and are relatively protected from hypothalamic inflammation^68^. Mechanistically, SFA can trigger a microglial inflammatory program characterized by pro-inflammatory cytokines production and oxidative stress^69–73^. Although brain lipidomic data in acute HFD paradigms (3-7 days) are limited, and generally reporting minimal or no changes in males^74,75^, we cannot exclude that subtle sex-dependent differences in lipid metabolism may shape microglial function in our model. Indeed, estrogens are potent regulators of brain lipid composition and metabolism^76^, and our data reveal sex-dependent differences in microglial lipid profile already under basal conditions, as well as divergent lipid remodeling in response to 3 days of HFD (Fig. S2C-E). In addition, sparse studies have shown that females have limited microglial morphological changes and sustain higher IL-10 expression under HFD^77,78^, consistent with a more protective phenotype mediated in part by ERα signaling^78,79^ and the CX3CR1–CX3CL1 axis, which dampens inflammation^77^. Importantly, microglia also harbor intrinsic sex-specific metabolic and transcriptional programs^80–84^ that persist independently of circulating gonadal hormones^85^. Taken together, these data support a model in which sex-specific lipid environments, hormone-dependent signaling, and cell-intrinsic microglial properties converge to generate divergent neuroinflammatory and metabolic responses to HFD.

Our data highlight a central role for antioxidant pathways in mediating the protective function of microglia in female mice fed an HFD. To date, sex-related differences in microglial bioenergetics has been largely underexplored, although some studies suggested sex-dependent regulation in pathophysiological contexts, such as aging or Alzheimer’s disease^86–88^. Previous work has documented rapid metabolic reprogramming, exclusively in male microglia, under acute HFD. Within 3d of HFD, ARH male microglia display enhanced mitochondrial respiration and ATP production in a UCP2-dependent manner, and deletion of microglial UCP2 attenuates inflammation and impairs fatty acid utilization, underscoring the importance of mitochondrial and lipid metabolism in early microglial adaptation to nutritional excess^66^. By contrast, in our experimental paradigm, male microglia did not display major mitochondrial remodeling over the first 3 days of HFD, and we did not detect cytokine induction (not shown). These discrepancies likely reflect differences in dietary composition (for both chow and HFD), experimental design, and/or circadian regulation of microglial immunometabolism^89^. Lipoprotein lipase (LPL) similarly regulates microglial responses to HFD in males by controlling phagocytic capacity and mitochondrial integrity, reinforcing the tight coupling between lipid metabolism and mitochondrial function^90^. Metabolic flux analyses indicate that during HFD, microglia preferentially oxidize fatty acids over glucose or glutamine, further comforting our results on their role in buffering lipid overload and preserving neuronal function^89^. Acute HFD drives profound metabolic remodeling in hippocampal microglia as well, including mitochondrial fission, inhibition of respiratory complex II, and production of succinate, itaconate, and lactate from palmitate β-oxidation, adaptations linked to enhanced hippocampal-dependent memory^36^. Recently, De Biase and colleagues reported that microglial TFAM deletion does not alter baseline microglial density^91^, a finding that we confirm in our study. Whereas they observed only subtle changes in microglial responses to acute inflammatory challenge (lipopolysaccharide injection), our data show that, under HFD conditions, intact mitochondrial function is essential^91^, further underscoring that caloric excess engages a noncanonical metabolic program in female microglia. Together, this evidence emphasizes the sex- and context-specific nature of microglial metabolic adaptation to nutrient excess.

Our findings demonstrate that HFD rapidly and selectively activates mTORC1 signaling in female but not male microglia, where it is instead reduced, providing the first evidence for a sex-specific signaling pathway operating in female microglia. Notably, this response contrasts sharply with that of male microglia, in which HFD predominantly engages inflammation-related pathways such as NF-κB^21^, highlighting a pronounced sexual dimorphism in microglial metabolic and signaling programs. This observation aligns with previous reports of sex differences in mTOR protein abundance, specifically lower expression in male microglia^92^, and extends them by demonstrating reduced mTORC1 pathway activation in male microglia. mTORC1 plays a central role in regulating immune cell function^93^, including that of microglia^29,94,95^, by integrating cytokine, nutrient, and hormonal signals to align cellular metabolic activity with available resources and the surrounding immune microenvironment^96^. Beyond this integrative role, mTORC1 acts as a potent pro-oxidant hub: it drives mitochondrial biogenesis and oxidative phosphorylation through PGC-1α, YY1, and 4E-BP1-dependent mechanisms^53,63^, thereby increasing electron transport chain flux and mitochondrial ROS production^55,97^, while concurrently restraining mitophagic clearance of damaged, ROS-generating mitochondria, since constitutive mTORC1 activation impairs PINK1/Parkin-dependent mitophagy and allows dysfunctional mitochondria to accumulate^98^. In myeloid cells, this pro-oxidant activity has direct functional consequences: mTOR signaling promotes NOS2 expression^99^, and mTOR-dependent oxidative stress licenses trained immune responses in monocytes exposed to oxidized lipids^100^. These observations position sustained mTORC1 activity as a plausible driver of the elevated ROS that we observe in female microglia after prolonged dietary exposure.

The relationship between mTORC1 signaling and redox homeostasis is nonetheless context-dependent. In settings of primary mitochondrial dysfunction, mTORC1 inhibition can instead lower ROS by restoring autophagic flux and mitochondrial quality control, and loss of mTOR complex 2 (mTORC2) increases mitochondrial ROS^101,102^, underscoring that the redox output of mTOR signaling depends on cellular context and on which mTOR complex is engaged. In our model, the convergence of mTORC1 activation, mitochondrial remodeling, and rising ROS strongly supports a pro-oxidant contribution of mTORC1 specifically within female microglia.

In conclusion, we show that in female mice exposed to caloric excess microglia, redox status emerges as a more critical regulatory axis than classical inflammatory activation, in contrast to what is typically observed in males. This raises the possibility that therapeutic strategies primarily targeting inflammation, largely developed and validated in male-biased models, may be suboptimal for women. More broadly, these insights have implications for a spectrum of pathological conditions in which microglia are central players, including neurodegenerative and neuropsychiatric disorders, for which obesity is a known risk factor^103–105^. Our findings therefore reinforce the need for sex-adapted therapeutic strategies and argue that biological sex should be incorporated as a fundamental design variable rather than a post-hoc consideration in treatment development.

### Limitations of the study

Several limitations of the present study warrant consideration. First, our loss-of-function approaches, including the Raptor, TFAM, and Hif1α KD, were microglia-specific but not spatially restricted, targeting microglia across the brain rather than selectively within the hypothalamus or ARH. As a result, we cannot exclude contributions to the observed *in vivo* phenotypes from microglia in other brain regions. Region-restricted, AAV-based approaches could help resolve this limitation, but they remain technically demanding and variable in efficiency, which currently limits their use for this purpose^106^. Second, the upstream signals driving the sex-specific microglial responses described remain to be identified, as the relative contributions of circulating nutrients, saturated fatty acids, sex hormones, and gut microbiota-derived signals to microglial activation in the female hypothalamus are still unknown.

## STAR METHODS

### Ethical considerations

All procedures involving live animals were approved and carried out in accordance with the National and European Directives 2013/63/EU, the French Ministry of Agriculture and Fisheries and the Ethical Committee of the University of Bordeaux and of the University of Lille for Animal Experimentation (authorizations #3959, #13394, #13395 and APAFIS#2617-2015110517317420 v5). Maximal efforts were made to avoid or reduce any suffering as well as to reduce the number of animals used.

### Animals

Male and female *C57BL/6J* (Janvier, France), *MitoTag (B6N.Cg-Gt(ROSA)26Sortm1(CAG-EGFP*)Thm/J)* (IMSR_JAX:032675), *tfam (tfam^fl/fl^)* (kindly provided by Dr A. Mourier, Bordeaux, France), *Raptor^fl/fl^ (B6.Cg-rptor^tm1.1Dmsa^/J)* (IMSR_JAX:013188), *Cx3cr1-EGFP*, and *Cx3cr1-creERT2 (B6.129P2(C)-Cx3cr1^tm2.1(cre/ERT)Jung^/J)* (IMSR_JAX:020940, Jackson Laboratory), *Hif1*_α_ *(hif1*_α_ *^fl/fl^)* (kindly provided by Dr Hamid-Reza Rezvani, Bordeaux, France), were used in this study. Male mice expressing tamoxifen-inducible Cre recombinase *(CreERT2)* in cells expressing *CX3CR1 (Cx3cr1-creERT2)* were crossed with *MitoTag* or *tfam^fl/fl^* or *Raptor^fl/fl^*to generate *CX3CR1*MitoTag*, *tfam* and *Raptor* mice and then crossed with female mice harboring conditional alleles of *MitoTag* or *tfam^fl/fl^* or *Raptor^fl/fl^*. Eight weeks old MG-MitoTag, MG-TFAM^KD^, MG-Raptor^KD^ mice and their littermate controls (WT fl/fl) were administered daily with tamoxifen by oral gavage at a dose of 150 mg/kg BW for 5 consecutive days. Experiments were performed 3 weeks after the last tamoxifen administration to allow the replacement of peripheral monocytes. All experiments were performed between 7 and 12 weeks of age at the onset of the experimental procedures.

Mice were single housed in standard plastic rodent cages under a 12:12 h reversed light/dark cycle (lights on at 3:00 h) at 22 ± 2°C. Cages and enrichment (cellulose nestlets, wooden sticks and cardboard tunnels) were changed fortnightly. Mice received a standard chow diet (Standard Rodent Diet A03, SAFE, France; 3.236 kcal/g; 13.5% calories from lipids, 25.2% calories from proteins and 61.3% calories from carbohydrates) and water *ad libitum*. For the acute or long-term high-fat diet exposure, mice were switched to a commercial high-fat diet (HFD) (D12492, Research Diets, USA; 5.24 kcal/g; 20% calories from proteins, 20% calories from carbohydrates, 60% calories from lipids) and were maintained on the diet for 1, 2 or 3 days (short-term exposure) or for 6 weeks (long-term exposure). Body weight and food intake was recorded daily for the short-term exposure to HFD and weekly for the long-term exposure to HFD, except during the first week after the diet switch, where it was recorded daily.

Body composition was assessed using nuclear echo magnetic resonance whole-body analysis on an EchoMRI 900 system (EchoMedical Systems, USA), following a previously validated protocol^107^. In brief, we recorded each animal’s weight and secured the mouse in a movement restrainer before inserting it into the EchoMRI to ensure accurate readings. We acquired all measurements in duplicate at a fixed time of day while animals had free access to food, then extracted fat mass and lean mass values for downstream analysis.

All animals were used in scientific experiments for the first time. This includes no previous exposures to the diet. Number of animals used in each experiment is indicated in the figure legends. Mice were allocated to experimental groups taking care to have similar body weight and fat mass content per group before the start of the experiments.

### Microglia Isolation from Adult Mouse Brains

For *ex vivo* experiments, microglia were isolated from adult mice hypothalamus or whole brain, as previously published^43^. Briefly, mice were sedated with Xylazine (20 mg/kg, Paxman, Virbac, France) and then euthanized with an overdose of pentobarbital (400 mg/kg, Euthasol Vet, Dechra, France) and then were transcardially perfused with ice-cold phosphate-buffered saline (PBS) to remove circulating blood. The hypothalamus was dissected, and microglia were isolated. For whole-brain microglia isolation, the olfactory bulbs and cerebellum were removed before tissue processing. The remaining isolation procedure was identical to that used for hypothalamic tissue, with reagent volumes adjusted according to the manufacturer’s instructions for the ABDK kit (Miltenyi Biotec). At the end of the isolation procedure, CD11b cells, consisting predominantly of microglia, were collected, washed in ice-cold Dulbecco’s phosphate-buffered saline (DPBS), and either processed immediately for downstream applications or centrifuged and stored as cell pellets at −80 °C until further use, as specified in the relevant method sections. Peritoneal macrophages, CD11b^+^ cells from liver or perigonadal adipose tissue were extracted with the same technique following vendor’s guidelines (MACS kits, Myltenyi).

### Metabolomics and lipidomics

Frozen pellets of hypothalamus microglia (8000 cells per tube) were treated with 100 uL of cold 5:3:2 methanol:acetonitrile:water then agitated 30 min at 4 degrees C for metabolite extraction. Insoluble material was pelleted by centrifugation (10,000 g, 10 min, 4 degrees C) then a 40 uL aliquot was removed from each tube, dried gently under vacuum, then resuspended in 20 uL of 0.1% formic acid for metabolomics analysis. After aliquot removal, the remaining material was treated with 50 uL of cold methanol then again agitated at 4 degrees C for 30 min to extract lipids. Insoluble material was pelleted using the conditions above then an aliquot of supernatant was dried under gentle vacuum and resuspended in half the volume of methanol. Metabolomics data was acquired in MS^1^ mode on a Thermo Vanquish UHPLC coupled to a Thermo Orbitrap Exploris 120 mass spectrometer using 5 uL injection volumes in negative and positive ion mode runs. Molecules were separated on a Waters Acquity BEH C18 column (2.1 x 30 mm, 1.7 um) using a 5 minute reverse phase gradient at 450 μL/min. The negative polarity gradient utilized mobile phases: A = water, 10 mM ammonium acetate; B = 50% acetonitrile, 50% methanol, 10 mM ammonium acetate. Solvent gradient: 0-0.5 min 0% B, 0.5-1.1 min 0-100% B, 1.1-2.75 min hold at 100% B, 2.75-3 min 100-0% B, 3-5 min hold at 0% B. Positive mode utilized mobile phases: A = water, 0.1% formic acid; B = acetonitrile, 0.1% formic acid) and solvent gradient: 0-0.5 min 5% B, 0.5-1.1 min 5-95% B, 1.1-2.75 min hold at 95% B, 2.75-3 min 95-5% B, 3-5 min hold at 5% B. Eluate was introduced to the MS via electrospary ionization and data was acquired across the scan range 65-975 m/z. Lipidomics data was acquired on a Thermo Vanquish UHPLC coupled to a Thermo Q Exactive mass spectrometer using a 5 min C18 gradient exactly as previously described^108,109^. Metabolomics raw data files were converted to mzXML format using RawConverter; peaks were annotated and integrated using Maven alongside an in-house compound library. Lipidomics raw files were mined using LipidSearch (Thermo). Data for each omics was normalized to median and autoscaled in MetaboAnalyst prior to statistical analysis.

### PCR for mouse genotyping and *rptor* gene excision quantification

PCR on tail biopsies was performed as in^110^ by using specific primers in two reactions, to detect the presence of floxed *rptor* alleles (Forward: 5’-CTCAGTAGTGGTATGTGCTCAG - 3’; Reverse: 5’-GGGTACAGTATGTCAGCACAG – 3’; Excised Forward: TCCCATGCCTTTAAACCCCC) and Cre recombinase gene (two sets of primers to run together, wild-type primers Forward: 5’–AGCTCACGACTGCCTTCTTC–3’; Reverse: 5’– ACGCCCAGACTAATGGTGAC –3’; Cre primers, Forward: 5’-CGGCATGGTGCAAGTTGAATA-3’, Reverse: 5’-GCGATCGCTATTTTCCATGAG-3’).

To obtain the absolute quantification of *rptor* gene excision, we developed a protocol based. Genomic DNA was extracted following the manufacture’s guidelines (RNeasy and all prep DNA/RNA minikit, Qiagen). Then, digital PCR was run with Bio-Rad T100 thermocycler, Automated Droplet Generator and QX200 Droplet Reader (Bio-Rad). PCR mix contained a minimal amount of 80ng DNA, the Taq QX200 ddPCR EvaGreen Supermix (Bio-Rad, #1864034) and the following sets of primers (WT, Forward 5’-GATGGCCTTGTCCCTCTGAC-3’, Reverse 5’-GACAGGAGCAAGAGGTACGG-3’; Lox Forward 5’-TCCTGGTCACAACAGAGTGC-3’, Reverse 5’-GATCCAAGGGAAAGAGCCCC-3’; KO Forward 5’-GGGCGCAGTGAGTACTGTT, Reverse 5’-GTGGGCATCTCACAAAGGGT-3’).

PCR cycling parameters and PCR mix calculations are available upon request.

### Quantification of ROS production by flow cytometry

Intracellular ROS levels were assessed using CellROX™ Deep Red reagent (Molecular Probes) as a marker of oxidative stress. CellROX™ Deep Red is a cell-permeant dye that is non-fluorescent in its reduced state and exhibits bright near-infrared fluorescence upon oxidation by ROS. Whole brain microglial cells isolated by magnetic cell separation (MACS) were resuspended in 100 µL of 1XHank’s Balanced Salt Solution (HBSS) and incubated with 2.1 µM CellROX™ Deep Red reagent at 37 °C for 30 minutes. Fifteen minutes before the end of the incubation, cells were treated with 1 µM SYTOX™ Blue to label dead cells. Positive and negative controls were included in each experiment. For the positive control, cells were treated with 1 mM tert-butyl hydroperoxide (TBHP) for 30 min prior to CellROX™ staining to induce ROS production, whereas untreated cells served as the negative control. Following incubation, cells were analyzed using a BD FACS Canto™ II flow cytometer (BD Biosciences). CellROX™ Deep Red fluorescence was detected using 640 nm excitation and 660 nm emission, whereas SYTOX™ Blue was detected using 405 nm excitation and 450 nm emission. Live cells were identified based on the absence of SYTOX™ Blue staining, followed by singlet discrimination using forward- and side-scatter parameters. Between 20,000 and 30,000 events were acquired per sample. The percentage of CellROX™-positive cells and the mean fluorescence intensity (MFI) were quantified within the viable microglial population.

### Determination of total glutathione

Whole brain microglial cells isolated by MACS were lysed with 1% (wt/vol) sulfosalicylic acid and centrifuged at 13,000*g* for 5 min at 4 °C, and the supernatants were used for determining total glutathione (GSH concentration plus twice the concentration of GSSG), by using GSSG (0–50 μM) as a standard. Total glutathione was measured in reaction buffer (0.1 mM NaHPO_4_, 1 mM EDTA, 0.3 mM DTNB, 0.4 mM NADPH and glutathione reductase at 1 U ml^−1^, pH 7.5) by recording the increase in absorbance at 405 nm after the reaction of GSH with DTNB for 2.5 min at 15-s intervals using a Varioskan Flash reader (Thermo Fisher). Results are expressed as mean ± s.e.m. (nmol x 10^−5^ cells).

### Oroboros O2k respirometry

Mitochondrial respiration was assessed on permeabilized microglia using a high-resolution O2k oxygraph (Oroboros Instruments) (detailed STAR protocol, Kyriakidou, Duranthon, Monsorno, 2026). Isolated microglia, previously resuspended in respiration buffer containing in mmol / L: (120 sucrose, 50 KCl, 20 Tris 4 KH_2_PO_4_, 2 MgCl_2_, 2.5 glucose, pH 7.2 with HCl), were loaded into the chambers, which were left open for a few minutes to allow entrapped air bubbles to escape before sealing with the stoppers. After closure, the absence of air bubbles was verified, as bubbles can interfere with oxygen flux measurements. Oxygen consumption was continuously recorded using DatLab software. Following sample loading and after each reagent addition, oxygen flux was allowed to reach a stable steady state before subsequent measurements were performed, and transient fluctuations corresponding to loading or injection artifacts were excluded from the analysis.

To permeabilize microglia the optimal digitonin concentration (0.8 µg/ml) was determined empirically according to cell number to ensure plasma membrane permeabilization while preserving mitochondrial integrity. Complex I-supported respiration was measured following the addition of pyruvate (10 mmol /L), glutamate (10 mmol /L), malate (5 mmol /L), and ADP (2.5 mmol /L). Maximal electron transfer system (ETS) capacity was determined by stepwise titration of the uncoupler carbonyl cyanide m-chlorophenyl hydrazine (CCCP) by step of 0.1 µmol /L until maximal oxygen consumption was reached. The highest stable oxygen consumption rate obtained after uncoupler titration was recorded as maximal ETS capacity.

At the end of the experiment, cells were recovered from the chambers and pelleted by centrifugation at 400 × g or 16,000 × g, respectively, for 5 minutes before further biochemical analyses, when required. The O2k chambers were subsequently cleaned according to the manufacturer’s recommendations using sequential washes with Milli-Q water, 70% ethanol, and 100% ethanol before storage. Oxygen flux values corresponding to each respiratory state were extracted from DatLab software using the Flux Flow analysis function after exclusion of non-steady-state regions.

### Real-time qPCR (RT-qPCR)

qPCR was carried out in isolated CD11b^+^ and CD11b^−^ cells from liver, adipose tissue, peritoneal cells, whole brain or from hypothalami from chow-fed and HFD-fed, C57BL/6J, MG-Raptor^KD^, MG-TFAM^KD^ mice, either after short-term or long-term exposure to the diet switch.

Isolated cells were lysed in BL Buffer and RNA from lysed cells was isolated using the ReliaPREP^TM^ RNA Cell Miniprep (Promega). RNA was processed and analyzed according to an adaptation of published methods^112^. cDNA was synthesized from total RNA using Maxima Reverse Transcriptase (Thermo Scientific) and primed with oligo-dT primers (Fisher Scientific) and random primers (Fisher Scientific). qPCR was performed using a LightCycler 480 Real-Time PCR System (Roche, France). qPCR reactions were done in duplicate for each sample using transcript-specific primers, cDNA (4 ng) and LightCycler 480 SYBR Green I Master (Roche) in a final volume of 10 μL. Primers were designed using PerlPrimer. For the determination of the reference genes, the RefFinder method was used^113^ and depending on tissue, different reference genes were used. All primers are reported in table X. The relative level of expression was calculated with the comparative (2^−ΔΔCT^) method^114^.

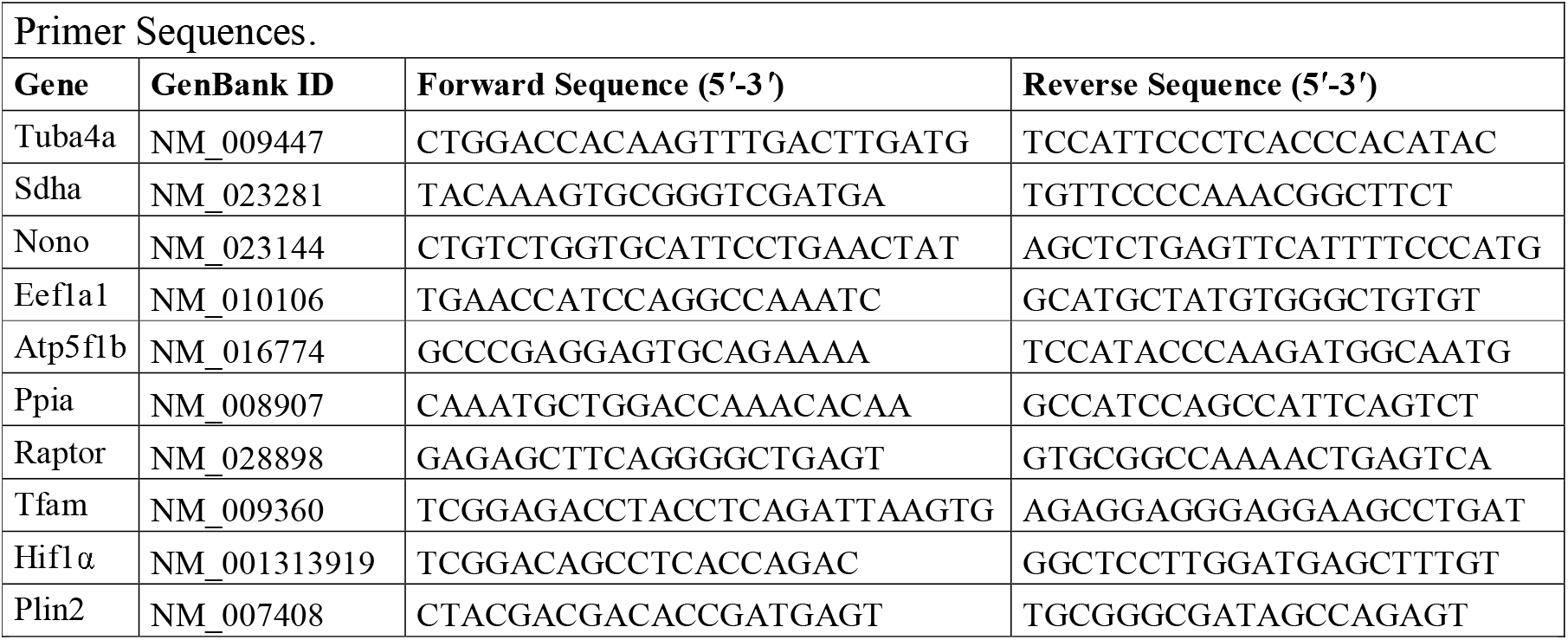

### Single-cell FACS sorting

For scRNAseq experiments, freshly dissociated single-cell suspensions from the hypothalamus of female and male C57BL/6 mice after 3 days of HFD, alongside chow-fed controls were sorted based on two markers: BV711-Cd45^+^ (Biolegend, clone 30-F11, catalog #103147) and APC.Cy7-Cd11b^+^(BD Pharmingen™; Catalog No:557657). Simple positive cells (CD45^+^, CD11b^−^ and CD45^−^ CD11b^+^) and double positive cells (CD45^+^CD11b^+^) are collected in order to collect all immune cell. Viable cells were selected based on side scatter gates, doublet exclusion and SYTOX blue viability dye 1:5000 (Fisher sci.). Cells were sorted using FACS Aria II upgrade at 4°C in pre-coated (2 hours at 37°C with PBS 10% FCS) 1.5 mL low binding tubes (Eppendorf) containing PBS-0.04% pure BSA (Fisher scientific, catalog 10743447) and counted with trypan blue dye before being processed.

### Single cell RNA sequencing experiment

Freshly isolated FACS-enriched CD45^+^, CD11b^−^, CD45^−^CD11b^+^ and CD45^+^Cd11b^+^ immune cells from hypothalamus of female C57BL/6 mice after 3 days of HFD or chow-diet were pooled using barcoding lipid technologies (10X Genomics, CG000391) as well for immune cells from hypothalamus of male C57BL/6 mice after 3 days of HFD or chow-diet. Each pooled contains a multiplexed of two samples, first, a mix of hypothalamus of 3 mice after 3-days HFD and a mix of hypothalamus of 3 mice after chow-diet. 27,000 cells/pool (corresponding to _≈_ 15,000 viable cells/sample) were loaded on the 10X chip to form Gel Bead-in-Emulsion in the Chromium Controller. Single-cell libraries were generated using the Single Cell 3’ reagent Kit v3.1 (10X Genomics, CG000388) as per the manufacturer’s protocol. cDNA was amplified by 14 PCR cycles for females, 15 PCR cycles for males were also performed for library preparation (single index PCR) and 6 cycles for CMO library. Libraries were pooled and sequenced on a NovaSeq PE150.

### Single cell RNA sequencing data processing

Fastq files were demultiplexed into individual samples and mapped to the mm10 reference genome (GRCm38-2020 Mus musculus) using the Cell Ranger multi pipeline, run on the MCIA server (Mesocentre de Calcul Intensif Aquitain). A count matrix was generated and converted into an RDS file for subsequent analysis in Seurat. Cells were filtered based on read counts, feature counts, and mitochondrial gene content for each sample. Doublets were identified by running DoubletFinder, scDblFinder, and scds, and cells identified as doublets at least two of the tools were removed. The samples were normalised by SCTransform, then integrated using CCA.

### Downstream analysis of microglial subclusters

Downstream analyses were performed in R v4.5.1. Clusters 0-3 were identified as microglial populations based on the expression of canonical microglial markers and automated cell-type annotation using SingleR v2.12.0 (main labels) with the ImmGen reference dataset obtained through celldex v1.20.0. Cells assigned to these clusters were extracted from the processed Seurat object (Seurat v5.3.1). The relative proportion of each microglial subcluster was calculated for each experimental condition as the number of cells assigned to that subcluster divided by the total number of microglial cells in the corresponding condition.

Pairwise differential expression between conditions was computed at the single-cell level using Seurat’s FindMarkers function with the Wilcoxon rank-sum test, after pooling clusters 0-3 and considering all detected genes. Genes were then ranked by the direction and significance of their differential expression (sign of the log_2_ fold-change × −log_10_ p-value, with infinite values capped at ±300), and this ranked list was used as input for gene set enrichment analysis with clusterProfiler::GSEA (v4.18.4) against the MSigDB Mouse Hallmark collection (msigdbr v25.1.1, category H; gene set sizes 50-500, eps = 0). P-values were adjusted using the Benjamini-Hochberg method, and gene sets with adjusted p<0.05 were considered significantly enriched.

Per-cell lipid metabolism scores were computed with Seurat::AddModuleScore using the REACTOME_METABOLISM_OF_LIPIDS gene set (MSigDB v2026.1.Mm), which quantifies the average expression of the gene set corrected by a matched randomly-selected control set.

### Bulk RNA sequencing

Hypothalamic microglial cells isolated by MACS were lysed in BL Buffer with Thioglycerol (ReliaPREP^TM^ RNA Cell Miniprep System, Promega). RNA was extracted using the ReliaPREP^TM^ RNA Cell Miniprep System and following the manufacturer’s instructions. The total RNA concentration and quality was determined using a 2100 Bioanalyzer Instrument (Agilent).

mRNA sequencing libraries were prepared according to Takara’s protocols using the SMART-Seq mRNA LP kit (Takara Bio, 634768) or SMART-Seq mRNA LP with UMIs kit (Takara Bio, 634762), Unique Dual Index Kit (97–192) (Takara Bio, 634753) or Unique Dual Index Kit (193–288) (Takara Bio, 634754). Profiling libraries were assessed using a LabChip GX Touch HT Nucleic Acid Analyzer (Revvity), with HT DNA NGS 3K Reagent Kit (Revvity, CLS960013) onto HT DNA X-Mark Chip (Revvity, CLS144006). Then libraries were quantified by qPCR onto LightCycler 480 II Instrument (Roche) with NEBNext Library Quant Kit (NEB, E7630L). Finally, libraries were sequenced at the PGTB facility with the NextSeq 2000 P3 Reagents (100 cycles), (Illumina, 20040559) or the NextSeq 2000 P3 XLEAP-SBS Reagent Kit (100 Cycles) (Illumina, 20100990) using the NextSeq 2000 sequencing system (Illumina). Paired-end sequencing reads of 50 bp resulted in an average of 32 to 40 million paired-end reads by sample.

Using the Aquitaine Mesocenter Computing Machines (MCIA), a part of raw data (Fastq files) were applied to our bioinformatic pipeline. Data quality assessment was performed by FastQC and MultiQC, read trimming with Cutadapt and Fastp. Following this, STAR was utilized to map the trimmed reads to the reference genome and quantify the mapped read, and the alignment was analyzed by Qualimap. Another part of raw data (Fastq files) were analyzed with Cogent NGS Analysis Pipeline v2.0.1 (Takara Bio). Read trimming was performed with Cutadapt, STAR was utilized to map the trimmed reads to the reference genome mm39 (GRCm39 Mus Musculus, release 113), SAMtools and bedtools were used to calculate Unique Start Stop (USS) positions for UMI and Subread for Gene expression counting.

R package DESeq2 was used to normalize and perform differential expression analysis on the quantified reads, identifying differentially expressed genes with FDR-adjusted p-value < 0.1 and preparing the data for functional enrichment analysis. Ensembl BioMart was used for ID mapping and feature extraction with the genome assembly. Gene set enrichment analysis identified enriched gene sets from the normalized count data and clusterProfiler was used to perform functional enrichment against the GO databases. Pheatmap package was used to cluster genes by group.

### Isolation of Mitochondria from Brain Tissue and Immunoprecipitation of GFP-OMM– Positive Mitochondria

Following euthanasia by cervical dislocation, whole brains from MG-MitoTag mice were rapidly isolated and transferred to PBS. Brain tissue was cut into small pieces and homogenized on ice in 2 mL of isolation buffer (IB; 150 mM KCl, 0.5 mM EGTA, 10 mM Tris, pH 7.2) supplemented with 0.1% fatty acid–free BSA (Euromedex) using a Potter– Dounce homogenizer. The homogenate was centrifuged at 500 × g for 10 min at 4 °C to remove debris. The resulting supernatant was collected and centrifuged at 5,000 × g for 10 min at 4 °C to pellet the mitochondria-enriched fraction. The pellet was resuspended in Isolation buffer (IB) containing in mmol /L : 150 MCl, 0.5 EGTA, 10 Trizma, pH 7.2 with HCl) supplemented with protease inhibitors (cOmplete Mini, Roche). Dynabeads Protein G (1.5 mg; Invitrogen) were incubated with 5 µg anti-GFP antibody (IgG1κ, Roche) for 45 min at room temperature under gentle rotation (20 rpm) after three washes in PBS containing 0.02% Tween-20. Antibody-bound beads were washed by magnetic separation and resuspended in 200 µL IB supplemented with protease inhibitors. The mitochondria-enriched fraction was then added to the antibody-coupled beads (final volume ≥ 500 µL) and incubated for 2 h at 4 °C under gentle rotation. The unbound fraction (flow-through) was collected, and beads were washed six times with 200 µL IB. Following washing, bead-bound mitochondria were collected for downstream analysis. Bound mitochondria were eluted by incubating the beads in 1× LDS sample buffer (Bolt, Thermo Fisher Scientific) for 10 min at 60 °C. The elution fraction was collected and supplemented with 1% DTT (Sigma-Aldrich) and nuclease (1:100; Pierce Universal Nuclease for Cell Lysis, Thermo Scientific).

### Western-Blot on isolated Mitochondria

The elution fraction of isolated GFP-OMM–positive mitochondria was separated by SDS– PAGE on 4–12% acrylamide gels (Invitrogen) together with a PageRuler Plus Prestained Protein Ladder in MES SDS running buffer (Invitrogen) and subsequently transferred to 0.2 µm nitrocellulose membranes. Membranes were stained with Ponceau S (0.1% in 5% acetic acid) to verify protein transfer and then blocked for 1 h at room temperature in 2.5% (w/v) non-fat dry milk prepared in Tris-buffered saline (20 mM Tris, 150 mM NaCl, pH 8) containing 0.05% Tween-20 (TBST). Membranes were incubated overnight at 4°C with OXPHOS cocktail primary antibodies (Abcam) diluted 1:1000 in milk-TBS. After washing in TBS, membranes were incubated for 1 h at room temperature with CyDye 800 goat anti-mouse secondary antibodies (Amersham) diluted 1:2000 in TBS. Fluorescence signals were detected using an ImageQuant 800 imaging system and band intensities were quantified with Fiji. Protein levels were normalized to the Ponceau S staining signal.

### Kinase profiling

Kinase activity profiles of isolated hypothalamic microglial cells were determined via PamChip^®^ peptide microarrays on a PamStation 12 (PamDx International B.V., a company formerly known as PamGene, Netherlands, https://pamdx.com/). PamChip^®^ microarrays contain immobilized 12-15 amino acids peptides derived from known or putative phosphorylation sites. These peptides get phosphorylated depending on the kinase activity in the lysates. Phosphorylated peptides were detected using assay-specific antibody-based fluorescence detection, and fluorescence signals were recorded with a charge-coupled device (CCD) camera.

Hypothalamic microglial cells were washed three times with DPBS after MACS to wash out BSA and be able to quantify the protein concentration of the sample. Then, cells were lysed with RIPA Buffer (NaCl 150 mM, tris-base 50 mM, 541 NP40 1% w/v, SDS 0.1% w/v, deoxycholate 0.5% w/v and EDTA 5 mM, pH 8) supplemented with protease inhibitor cocktail (Sigma-Aldrich) and Halt^TM^ phosphatase inhibitor cocktail (ThermoScientific). Protein concentration was determined using the Pierce^TM^ Micro BCA^TM^ Protein Assay Kit (Thermo Scientific, #23235) according to manufacturer’s instructions.

For kinase activity profiling, Protein-Tyrosine Kinase (PTK) and Serine/Threonine Kinase (STK) chips containing 196 and 140 peptides respectively, targeting the main kinase families were used. All reagents were supplied by PamDx. For PTK activity profiling, 3 µg of total protein amount was dissolved in 4 µL 10× PK buffer, 0.4 µL 100× BSA, 4 µL 4 mM ATP, 0.6 µL FITC conjugated antibody, 0.4 µL 1M DTT, 4 µL 10× PTK additive and filled up with distilled water to 40 µL total volume and then loaded on the chip. Prior to sample loading, a blocking step was performed using 30 µL of 2% BSA. For STK activity profiling, 0,5 - 2 µg of total protein amount was dissolved in 4 µL 10× PK buffer, 0.4 µL 100× BSA, 4 µL 4 mM ATP, 0.5 µL STK primary Antibody mix and filled up with distilled water to a total volume of 40 µL and then loaded on the chip. Prior to sample loading a blocking step was performed loading 30 µL of 2% BSA. In the second part of the STK program, each chip was loaded with the detection mix (30 µL, consisting in 0.4 µL FITC conjugated antibodies dissolved in 3 µL 10× AB buffer and 26.6 µL water). To determine the kinetic of kinase activity, the samples were pumped several times (cycles) through the microarray and imaged at certain cycle passages. For all experiments, 3-5 biological replicates were used.

Signal intensities were analyzed in BioNavigator software (PamDx) and expressed as log fold change. Prediction of kinases responsible for altered phosphorylation between conditions was analyzed using the upstream kinase analysis app (2018 version; PamDx) and was based on multiple kinase–substrate relationship databases.

### Multi-Electrode Array (MEA) experiments

Control and MG-Raptor^KD^ mice brains were sectioned at 300 µm thickness and used for MEA recordings. After 1 hr of recovery in aCSF (identical composition to aCSF used for imaging) at room temperature, slices were placed onto the MEA chip. Slices were precisely placed relative to the electrode array under visual guidance using a microscope. To stabilize the slice in the recording chamber, a thin plastic ring and a metal harp were used. The slice was acclimatised in prewarmed oxygenated aCSF for at least 15 minutes before the recordings began. The recording chamber was maintained at 32°C. MEA recordings were performed for 30 minutes (1 slice per genotype) under continuous perfusion of prewarmed aCSF. The data was analysed offline using the “Multi channel analyser” software. Signals were filtered using Butterworth low pass filter with 100 Hz cutoff. Spikes were extracted from the raw data using the ‘spike detector’ tool. We applied a manual threshold (falling edge of −15µV), which was uniformly applied to all recording channels. Spikes were counted for the selected electrodes/channels (in the hypothalamus) using the ‘spike analyzer’ tool and plotted using GraphPad Prism.

### Transmission electron microscopy sample preparation and imaging

Freshly dissected mediobasal hypothalami (containing the ARH) were initially fixed in 2.5% glutaraldehyde diluted in phosphate buffer 0.1M pH 7.4 for 4 hours at 4°C. Following three rinses in phosphate buffer 0.1M pH 7.4, samples were post-fixed overnight at 0°C in a solution containing 1% osmium tetroxide (OsO) and 15 mg/ml potassium ferrocyanide (K Fe(CN)) in phosphate buffer 0.1M pH 7.4. After post-fixation, tissues were washed again with phosphate buffer 0.1M pH 7.4 and subsequently dehydrated through a graded acetone series. Samples were then progressively embedded in Epon resin. Ultrathin sections (80 nm) were obtained, contrasted with lead citrate, and examined at an accelerating voltage of 80 kV using a Hitachi 7650 transmission electron microscope (Electron imaging facility of Bordeaux Imaging center). Quantitative analysis of the acquired images was performed using Fiji software.

### Two-photon imaging

CX3CR1-EGFP Mice were anesthetized with isoflurane then cervical dislocation was performed before decapitation. Brains were quickly extracted and placed in ice-cold NMDG based solution containing (in mM) 1.25 NaH_2_PO_4_, 2. 5 KCl, 7 MgCl_2_, 20 HEPES, 0.5 CaCl_2_, 28 NaHCO_3_, 8 D-glucose, 5 L(+)-ascorbate, 3 Na-pyruvate, 2 thiourea, 93 NMDG and 93 HCl 37%; pH: 7.3−7.4; osmolarity: 305−310 mOsM. Brain slices (coronal) of 300µm thickness were prepared by gluing the extracted brains (cerebellum up and olfactory side down) dorsal side facing the blade. Mediobasal hypothalamic sections were collected (consisting of third ventricle with Median eminence and arcuate nucleus). Slices were left to recover for 15 min in NMDG at 34°C and for at least 45 min at room temperature in aCSF containing (in mM) 124 NaCl, 2.5 KCl, 1.25 NaH_2_PO_4_, 2 MgCl_2_, 2.5 CaCl_2_, 2.5 D-glucose and 25 NaHCO_3_; pH: 7.3−7.4; osmolarity: 305−310 mOsM.

For imaging GFP-labeled microglial cells in acute brain slices a commercial 2-photon microscope (Prairie Technologies) was used. Its wavelength was tuned to 920 nm. 4D imaging (a z-stack of 30 µm thickness with a step size of 1 µm, acquired every 15 minutes) was performed using a 40X water immersion objective with an NA of 1.0 (Plan Apochromat, Zeiss). Laser power was between 20-40 mW in the focal plane. The fluorescence signal was collected by PMT detectors. Images were acquired with a pixel size of 576 nm over a 295×295 μm^2^ field of view and pixel dwell-time of 12.4 μs. Image acquisition was controlled by Prairie View (v.4.0). Imaging was done under continuous perfusion of aCSF at the rate of 2 ml/min. The imaging chamber was maintained at 34°C.

### Brain tissue sampling

Mice were sedated with Xylazine (20 mg/kg, Paxman, Virbac, France) and then euthanized with an overdose of pentobarbital (400 mg/kg, Euthasol Vet, Dechra, France). Mice were transcardially perfused with ice-cold PBS pH 7.4, followed by 4% PFA. Brains were extracted and post-fixed in 4% PFA overnight at 4 °C, then cryoprotected in 30% sucrose in PBS at 4 °C. Coronal sections (30 μm) were cut using a cryostat (CM3050S, Leica), collected and stored in anti-freeze solution (30% ethylene glycol, 30% glycerol in KPBS) at −20 °C until further use.

### Histological analyses

Brains were sectioned coronally at 30 µm using a Leica cryostat. Free-floating brain sections were permeabilized in 0.5% Triton X-100 in PBS for 90 min at room temperature (RT), followed by incubation for 1 h at RT in blocking buffer consisting of 2% bovine serum albumin (BSA), normal goat serum (NGS), or normal donkey serum (NDS) prepared in permeabilization buffer, depending on the primary and secondary antibodies used. Primary antibodies were diluted in the corresponding blocking buffer and incubated with the sections overnight at 4 °C. The following primary antibodies were used: rabbit anti-Iba1 (1:1000; Wako, 019-19741), guinea pig anti-Perilipin 2 (PLIN2, 1:200; Progen, GP40), and goat anti-green fluorescent protein (GFP, 1:1000; Sicgen, AB0066200). The following day, sections were washed in PBS and incubated for 2 h at RT with the appropriate fluorescent secondary antibodies: goat anti-rabbit Alexa Fluor 647 (1:2000; Thermo Fisher Scientific, A21246), goat anti-guinea pig Alexa Fluor 488 (1:400; Thermo Fisher Scientific, A11073), donkey anti-rabbit Alexa Fluor 647 (1:1000; Jackson ImmunoResearch, 711-605-152), or donkey anti-goat Alexa Fluor 488 (1:2000; Thermo Fisher Scientific, A11055). Nuclei were counterstained with DAPI (1 µg/mL in PBS) for 5 min. Sections were then rinsed three times in 50 mM Tris-HCl (pH 7.5) and mounted with ProLong™ Gold Antifade Mountant (Invitrogen, P36930). Images were acquired using a Leica TCS SP8 confocal microscope (Leica Microsystems) equipped with ×20, ×40, or ×63 oil-immersion objectives and controlled with Leica Application Suite (version 3.5.7) at the Bordeaux Imaging Centre (BIC). Z-stack images were acquired with a z-step size of 0.3 µm.

### Image analysis

#### Image J

Dynamics of microglial processes was tracked in maximum intensity projections of the timelapse images, using ImageJ. The images of microglia from the first and last time point were overlaid. Using the *Segmented line* tool, individual process tips were traced in the overlay. Red trace indicates retraction and green indicates extension. To quantify average shape dynamics of each microglia ImageJ plugin *MotiQ* was used^115^. Briefly, *MotiQ* consists of three steps: (A) Single cell image generation using *MotiQ cropper*, (B) Image segmentation using *MotiQ thresholder* and (C) Image analysis using *MotiQ 2D analyzer*. *MotiQ* analyses the shape dynamics, returning the sum of the total extended and retracted area. This analysis shows how the shape of microglia changes over time, which was then averaged across time. Each dot in the graph corresponds to an individual microglial cell.

#### Fiji

For quantification of Iba1 cells in the ARH, maximum-intensity projections of confocal z-stacks were generated and used for all analyses. The region of interest (ROI) corresponding to the AR was manually delineated using the polygon selection tool in ImageJ. Images were thresholded and converted to binary masks before particle analysis was performed using the Analyze Particles function. Only particles entirely contained within the ROI were included in the analysis. Iba1 cell number was quantified for each ROI and expressed as cell density (cells/mm²).

#### Imaris

Microglial morphometric analyses, quantification of Plin2 microglia, and GFP particle analysis within microglia were performed using Imaris software (version 10, Bitplane). All individual microglial cells within each field of view were individually reconstructed and analyzed. Microglial morphology was reconstructed using the *Filament Tracer* module, from which the number of branch points and total filament length were extracted for each cell. For volumetric analyses, Iba1, Plin2, and GFP signals were quantified using 3D surface rendering of confocal z-stacks in their respective channels. Identical rendering parameters were applied across samples within each experiment to ensure consistency. To specifically quantify Plin2 and GFP signals within microglia, only particles located inside the Iba1 volume were considered. To this end, masked channels corresponding to “microglial Plin2” or “microglial GFP” were generated using the Imaris mask function, restricting Plin2 or GFP signals to the Iba1 microglial volume. To account for differences in microglial cell size, Plin2 and GFP volumes were normalized to the total Iba1 volume of each individual microglial cell. The same approach was applied for the quantification of Plin2 colocalizing with GFP in MG-MitoTag mice. Finally, the proportion of Plin2 microglia was determined by manual counting using maximum intensity projections. Microglial cells were classified as Plin2 when masked Plin2 signal was clearly detectable within the Iba1 microglial soma and/or processes. The percentage of Plin2 microglia was calculated as the number of Plin2 microglial cells divided by the total number of Iba1 microglia per field of view.

### Statistical analyses

Statistical analyses were performed using GraphPad Prism 8 (GraphPad Software, San Diego, CA, USA). All values are expressed as mean ± standard error (SEM). Equality of variances and normality of distributions were verified. The level of statistical significance between groups was determined, depending on the data, using unpaired t-test, Welch’s test, one sample t-test, Kruskal-Wallis test, two-way ANOVA on repeated measures analysis with Tukey’s multiple comparisons when needed. A p-value < 0.05 was considered statistically significant.

## SUPPLEMENTAL INFORMATION

**Figure S1. Acute HFD exposure does not induce major alterations in microglial morphology or motility.**

**(A–B)** Representative immunohistochemical images of the ARH showing Iba1-immunoreactive microglia and quantification of microglial density after 1 or 3 days of HFD exposure in male and female mice (females, green; males, blue) (n=5, one-way ANOVA).

**(C)** Representative three-dimensional filament reconstructions of ARH microglia from chow- and HFD-fed mice after 1 or 3 days of diet exposure.

**(D-E)** Quantification of microglial morphology, including the number of branch points and total process length, following 1 or 3 days of HFD exposure (n=4, one-way ANOVA).

**(F)** Quantification of microglial volume in the ARH of chow- and HFD-fed male and female mice (n=4, one-way ANOVA).

**(G)** Representative two-photon microscopy images illustrating ARH microglial process dynamics in HFD-fed mice after 1 day of diet exposure (‘Ext.’: extension, ‘Retr.’: retraction).

**(H)** Quantification of microglial process motility following 1 or 3 days of HFD exposure (n=4-11, one-way ANOVA).

**Figure S2. Acute HFD exposure affects microglial transcriptome and lipidome.**

**(A)** Per-cell Reactome lipid metabolism module score (Seurat::AddModuleScore), shown as violins with overlaid box plots (median and IQR), faceted by cluster.

**(B)** PLS-DA analysis of untargeted lipidomic profiles obtained from MACS-isolated hypothalamic microglia from chow- and HFD-fed male and female mice following 3 days of diet exposure.

**(C)** Heatmap showing the relative abundance of major lipid classes across experimental groups after 3 days of HFD exposure.

**(D-E)** Lipid Ontology (LION) enrichment analysis of lipidomic datasets from female (D) and male (E) microglia following 3 days of HFD exposure. Enriched lipid-associated pathways and functional annotations are displayed according to enrichment score and significance (n = 5 mice/group).

**(F)** Representative 3D reconstruction of confocal images of hypothalamic microglia immunostained for Iba1 and the lipid droplet marker Perilipin-2 (Plin2) in chow- and HFD-fed mice after 3 days of diet exposure.

**(G)** Quantification of Plin2-positive lipid droplet content in ARH microglia after 3 days or 6 weeks of HFD exposure, normalized to chow-fed controls, and percentage of Plin2-positive microglia (cells lacking detectable Plin2 particles were considered Plin2-negative). Data are represented as the average per animal (n = 5, unpaired t-test, *p<0.05).

**(H)** Experimental paradigm.

**(I)** Number of differentially expressed genes and proteins identified in female and male microglia following 3 days of HFD exposure.

**(J)** Venn diagram showing the overlap of HFD-responsive genes between female and male microglia.

**(K-L)** Gene ontology enrichment analysis of differentially expressed genes in female (K) and male (L) microglia following 3 days of HFD exposure. Female microglia preferentially enriched pathways related to nutrient sensing, metabolism, and lipid homeostasis, whereas male microglia displayed enrichment of immune-related pathways. Dot size represents the number of differentially expressed genes associated with each pathway, while color indicates the adjusted *p* value, with darker red corresponding to greater statistical significance.

**Figure S3. Acute HFD exposure recruits the mTORC1 pathway in female microglia**

**(A–B)** Top significantly altered protein tyrosine kinases (PTKs) and serine/threonine kinases (STKs) in female (A) and male (B) microglia following 3 days of HFD exposure.

**(C)** Representative image of the PCR amplicons for genotyping (floxed versus wild-type alleles) and excision control (band at 280 bp) of the *rptor* gene in the cortex, peritoneal macrophages, white adipose tissue (WAT) and liver from control (Ctrl) and MG-Raptor^KD^ mice, 7 weeks after recombination induction (n=3/genotype).

**(D)** Quantification of *rptor* gene excision by droplet digital qPCR in forebrain MACS CD11b+ (microglia) and CD11b-cells (n=2-5).

**(E)** Relative *rptor* mRNA expression in CD11b+ and CD11b-cells from the forebrain (n=5-6, two-way ANOVA, ****p<0.0001).

**(F)** Relative *rptor* mRNA expression in peritoneal macrophages and other circulating peritoneal cells (n=7-23), CD11b+ and CD11b-cells from white adipose tissue (WAT, n=11), and liver (n=11-12) isolated from control and MG-Raptor^KD^ mice. Expression levels were normalized to control mice (n=11-23, two-way ANOVA).

**Figure S4. Microglial mTORC1 orchestrates early kinase signaling responses to HFD**

**(A–B)** Kinome trees illustrating kinase activity profiles in control female microglia following 1 day of HFD exposure relative to chow-fed controls (A) and in MG-Raptor^KD^ microglia relative to HFD-fed control mice after 1 day of HFD exposure (B). Kinases are organized according to their phylogenetic families. Red nodes indicate increased kinase activity, whereas blue nodes indicate decreased kinase activity. Node size reflects the magnitude of the kinase activity change.

**(C–D)** Top significantly altered protein tyrosine kinases (PTKs) and serine/threonine kinases (STKs) in control female microglia following 1 day of HFD exposure compared with chow-fed controls (C) and in MG-Raptor^KD^ microglia compared with HFD-fed control mice after 1 day of HFD exposure (D). Bars represent median kinase activity scores. Color intensity reflects kinase specificity, with darker red indicating higher specificity scores.

**(E)** Gene ontology enrichment analysis of differentially expressed genes identified by bulk RNA sequencing in MG-Raptor^KD^ microglia compared with HFD-fed control microglia following 1 day of HFD exposure. Enriched pathways are primarily associated with cell cycle regulation and cellular organization. Dot size represents the number of differentially expressed genes associated with each pathway, whereas color indicates the adjusted *p* value.

**(F)** Microglial density (Iba1+ cells) in the ARH of MG-Raptor^KD^ mice exposed to HFD for 1d or 3d (n=5, unpaired t-test).

**(G)** Representative image of the multielectrode array (MEA) recording setup illustrating the hypothalamic nuclei analyzed, including the ARH, ventromedial hypothalamus (VMH), dorsomedial hypothalamus (DMH), lateral hypothalamus (LH), and median eminence (ME).

**(H)** Quantification of spontaneous neuronal firing activity in the indicated hypothalamic regions following chronic HFD exposure (n=4, two-way ANOVA, *p<0.05, ***p<0.001).

**(I)** Percentage body weight changes relative to baseline in MG-Raptor^KD^ male mice compared to Ctrl (n=20-32, two-way ANOVA RM, ns: non-significant).

**Figure S5. Evaluation of the role of HIF1_α_ and of mitochondria in the effect of microglial mTORC1 in the female response to calorie excess.**

**(A)** Relative *Hif1*_α_ mRNA expression in microglia isolated from female mice exposed to HFD for 1d or 6w, versus chow-fed controls (n=8-10, one-way ANOVA).

**(B)** Experimental design of the inducible microglia-specific *Hif1*_α_ knock down (MG-HIF1α^KD^) mouse model. Tamoxifen was administered to 8-week-old mice and experiments were performed 3 weeks later.

**(C)** Relative *Hif1*_α_ mRNA expression in microglia isolated from control and MG-HIF1α^KD^ mice (n=3, Welch’s test).

**(D)** Percentage body weight changes relative to baseline in MG-HIF1α^KD^ female mice compared to Ctrl during chronic HFD exposure (n=9-12, two-way ANOVA RM).

**(E)** Percentage body weight changes relative to baseline in MG-HIF1α^KD^ male mice compared to Ctrl during chronic HFD exposure (n=9, two-way ANOVA RM).

**(F)** Relative *tfam* mRNA expression in microglia isolated from control and MG-TFAM^KD^ mice (n=4-6, Welch’s test, **p<0.01).

**(G)** Oxygen consumption rates of permeabilized microglia isolated from control and MG-TFAM^KD^ mice during the different respiratory states of the SUIT protocol. Values are expressed as oxygen flux (JO; amol O ·min ¹·cell ¹) (n=6-9, two-way ANOVA RM, **p<0.01, ***p<0.001, ****p<0.0001).

**(H)** Quantification of microglial density (Iba1+ cells) in control and MG-TFAM^KD^ mice (n=5, unpaired t-test).

**(I)** Percentage body weight changes relative to baseline in MG-TFAM^KD^ mice male mice compared to Ctrl (n=10-18, two-way ANOVA RM, ns: non-significant).

## ACKNOWLEDGEMENTS

A.N. was supported by the Institut Universitaire de France (IUF), the University of Bordeaux, and the National Research Agency (ANR-23-CE14-0076 MicroNRJ; ANR-24-NEU2-0009 EMPATHY), the French government through the University of Bordeaux’s IdEx “Investments for the Future” program / GPR BRAIN_2030, the Fondation pour la Recherche sur le Cerveau (FRC), and the Groupe Lipides et Nutrition (GLN). A.N., G.C. and E.K. received funding from the European Union’s Horizon research and innovation program under MSCA Doctoral Networks 2021, grant No. 101072759 (ETERNITY, FuEl ThE bRaiN In healtThY aging and age-related diseases). D.C. acknowledges support from INSERM, Nouvelle-Aquitaine Region, Agence Nationale de la Recherche (ANR-18-CE14-0029; ANR-21-CE14-0018, ANR-22-CE14-0016, ANR-23-CE14-0037, ANR-25-CE14-7688), University of Bordeaux’s IdEx ‘‘Investments for the Future’’ program/GPR BRAIN_2030, and Fondation pour la Recherche Medicale (FRM EQU202303016291). C.A. acknowledges support from the University of Bordeaux and FRM (FRM-ARF201809006962). The Biochemistry and Biophysics Platform (BioProt) of the Bordeaux Neurocampus, equipped with the PamStation (PamDx analysis), is supported in part by the GPR BRAIN_2030. JPB is funded by the European Research Council (ERC) Advanced Grant NeuroSTARS (101199747).

This work benefited from the support and help from the PUMA Tissue and Cell Isolation Facility (M. Maitre and H. Doat), the Genotyping Facility (E. Huc, C. Gatuingt-Chasseriaud, D. Gonzalez, G. Laplagne) and the Animal Facility (R. Racunica and F. Corailler) at the NeuroCentre Magendie, Inserm U1215. Part of the experiments (mRNA sequencing) were performed at the PGTB (https://doi.org/10.15454/1.5572396583599417E12) with the help of Préscillia Alves-Gomes and Zoé Compagnie. The microscopy and the image analysis were done in the Bordeaux Imaging Center a service unit of the CNRS-INSERM and Bordeaux University, member of the national infrastructure France BioImaging supported by the French National Research Agency (ANR-24-INBS-0005 FBI BIOGEN). The help of Monica Fernandez Monreal and Sebastien Marais is acknowledged. We thank Atika Zouine, Vincent Pitard and Jean-Michel Griffon for technical assistance at the Flow cytometry facility, UAR 3427, INSERM US 05, Univ. Bordeaux, F-33000 Bordeaux, France and Amy Argabright and Colin Anderson at the University of Colorado Metabolomics Facility.

## AUTHOR CONTRIBUTIONS

A.N., D.C., and C.A. conceived the studies, oversaw their design and execution, and wrote the final manuscript. C.A., E.K. and G.C. designed and carried out the experiments, analyzed the data, and generated figures. A.N. prepared the final figures. T.C., E.A., J.V., E.G., A.C., L.C., E.C., E.P., V.S., P.Z., S.J., and N.D. assisted with key experiments. S.L. ran the scRNAseq experiments, and O.K., G.A., C.A., and R.R. helped with bioinformatic analyses of the dataset. T.L.L. and F.M. performed transcriptomic experiments, while A.B. contributed to RNAseq bioinformatic analyses. P.G., U.V. N. and L.G. performed the 2-photon and MEA experiments. A.D.A. and J.A.R. performed lipidomics and metabolomics studies and A.D.A helped with interpretation of the data. M.D. and J.E. contributed to the kinomics experiments and analyses. G.M. contributed to interpretation of mitochondrial data. B.S. performed electron microscopy experiments. J.P.B. and D.J.B. assisted with glutathione measure. G.D. and A.M. conducted respirometry experiments and Western blot analysis of OXPHOS complexes. A.M. provided TFAM-floxed mice. All authors edited and approved the final manuscript.

## DECLARATION OF GENERATIVE AI AND AI-ASSISTED TECHNOLOGIES IN THE MANUSCRIPT PREPARATION PROCESS

During the preparation of this manuscript, the authors used Claude (Anthropic) to check grammar and improve the clarity of the English text. The authors reviewed and edited all output and take full responsibility for the content of the publication.

