## Supplementary figures and images for "A Female-Specific Microglial Redox Program Gates Susceptibility to Obesity"

### Figure S1

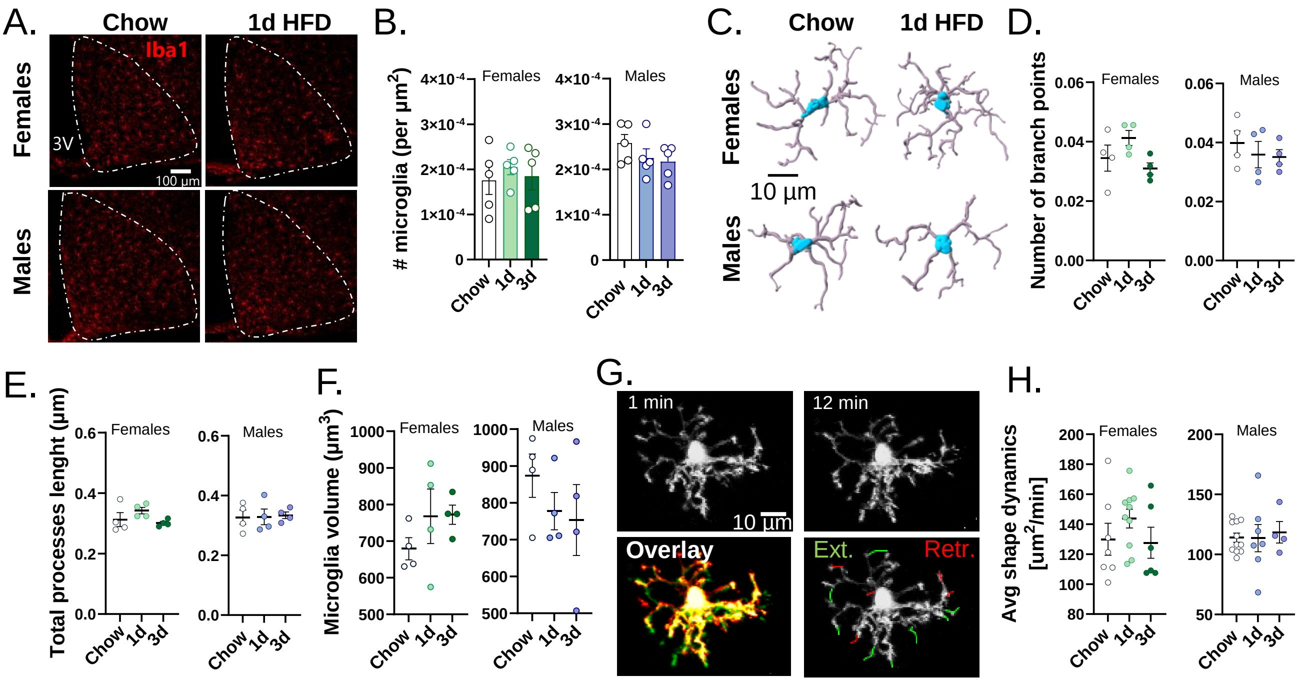

### Figure S2

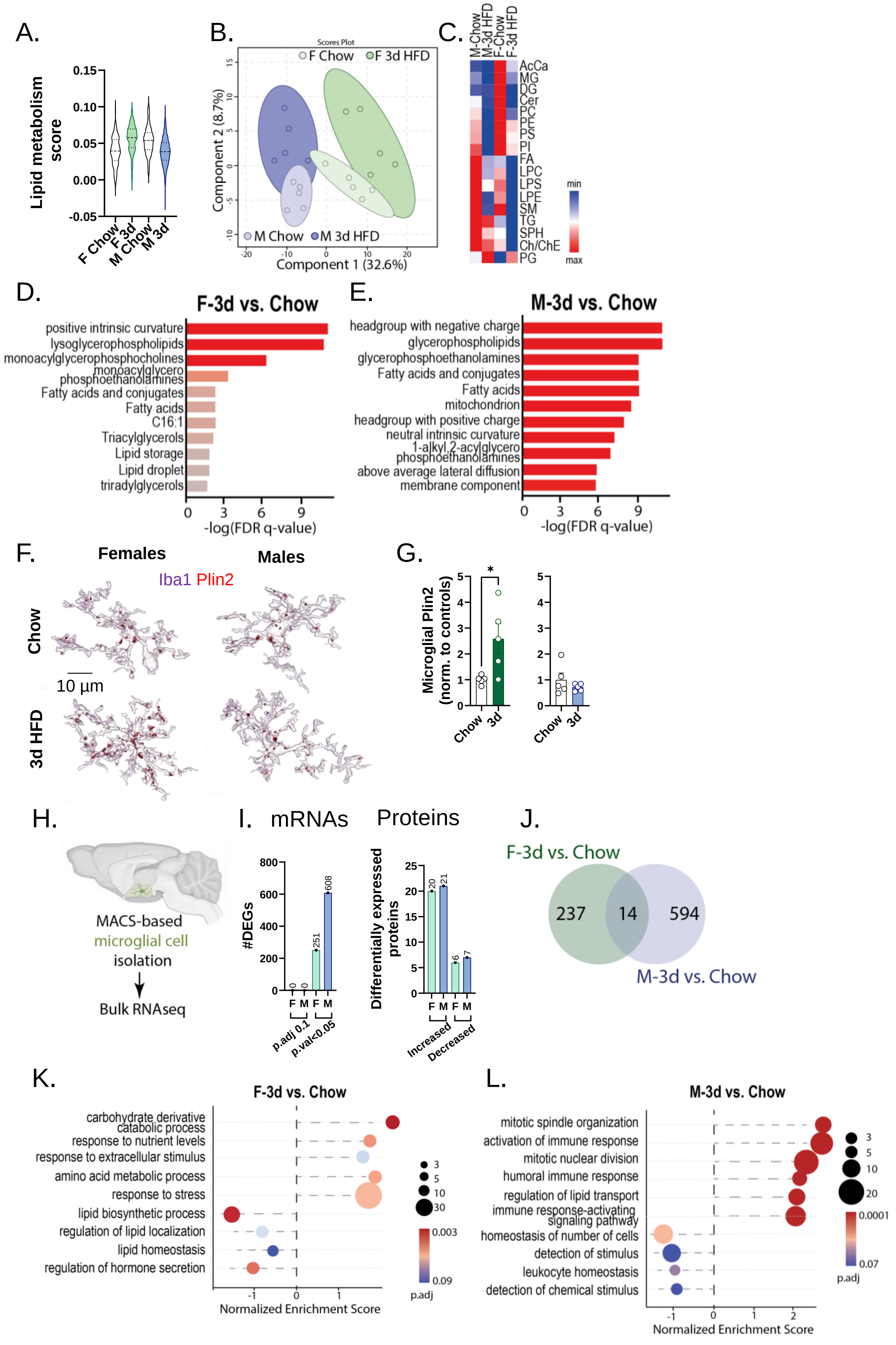

### Figure S3

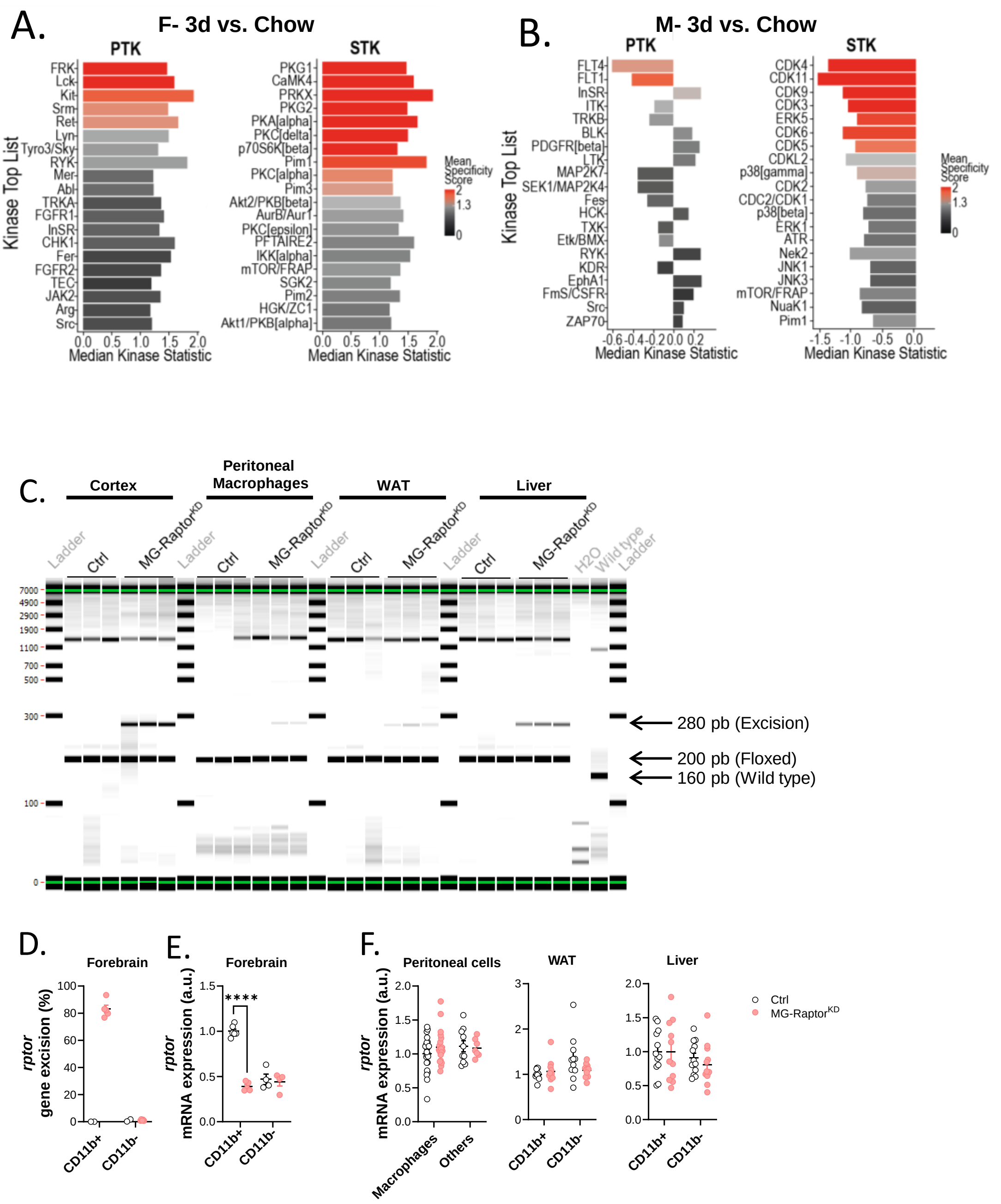

### Figure S4

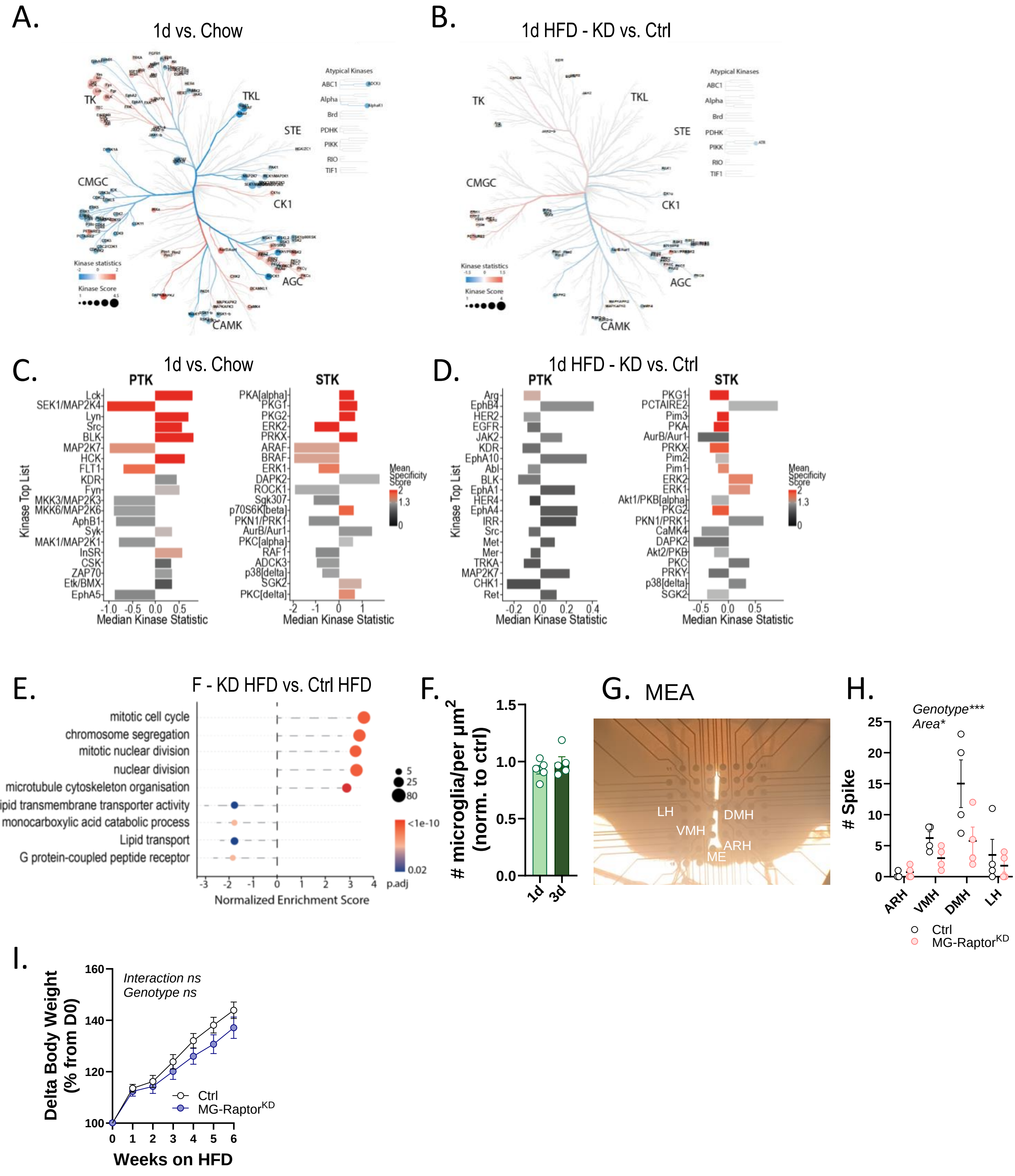

### Figure S5

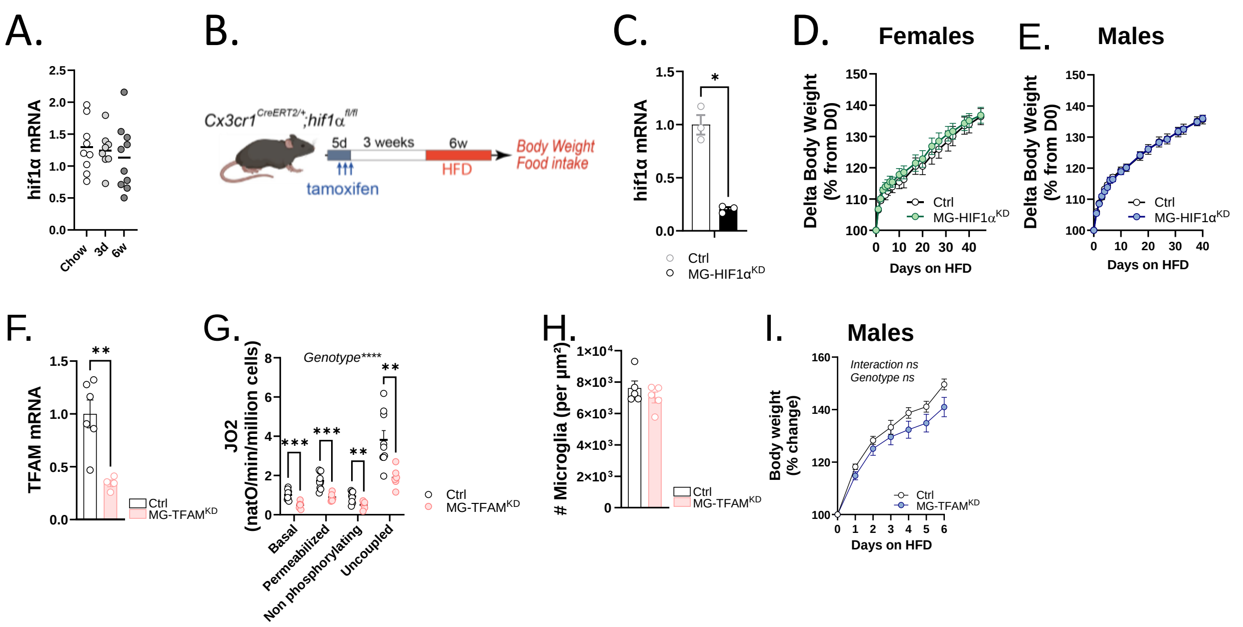
